# High resolution spatial transcriptomics decodes the microenvironmental determinants of response to Nr-CWS therapy in cervical precancerous lesions

**DOI:** 10.64898/2026.06.02.729727

**Authors:** Bei Wei, Qing Sun, Yingao Ma, Yunfeng Sun, Junzhe Li, Ye Lu, Xiaokang Li, Yuying Lan, Jianda Ma, Miao Ding, Tengfei Long

## Abstract

Immunotherapy with Nocardia rubra cell wall skeleton (Nr-CWS) can clear human papillomavirus (HPV) and induce regression of cervical precancerous lesions, yet many patients do not respond. The cellular and molecular basis for this heterogeneous clinical outcome remains unknown. Using high-resolution spatial transcriptomics, we profiled squamous intraepithelial lesions (SIL) from patients stratified by their subsequent response to Nr-CWS therapy. We discovered that non-responders were characterized by an immunosuppressive epithelial phenotype, defined by aberrant expression of the pro-inflammatory alarmin S100A9 in squamous epithelial cells (SECs), which was associated with poorer overall survival in cervical cancer. Non-responding lesions were enriched in proliferative (MKI67+) and pro-angiogenic (VEGFA+) SEC subpopulations and spatially organized into a pro-tumorigenic cellular neighborhood (CN-10) overexpressing S100A9 and ANXA1. In contrast, responders exhibited protective glandular-cell–enriched neighborhoods (CN-13) and supportive spatial interactions between lymphatic endothelial cells and fibroblasts. The unfavorable microenvironment was further defined by an immunosuppressive cloak of VCAN+ stromal cells and CD163+ M2 macrophages encircling SECs. Thus, response to Nr-CWS is determined by a pre-existing spatial network of epithelial-stromal-immune interactions, providing a roadmap for patient stratification and rational combination therapies.

## INTRODUCTION

Cervical cancer continues to be the most common gynecological malignancy globally, posing a significant threat to women’s health, particularly in low- and middle-income countries^1^. Persistent infection with high-risk human papillomavirus (HPV) has been established as the principal causative agent, driving the progression from precancerous cervical lesions to invasive carcinoma^2^. Although prophylactic HPV vaccines have demonstrated success in reducing the incidence of HPV-related diseases, they offer limited therapeutic benefit to individuals already infected. Therefore, promoting viral clearance and treating existing cervical lesions remain crucial strategies for preventing cervical carcinogenesis^3^.

The pathogenesis of HPV-induced cervical lesions and cancer is not yet fully understood. A key challenge lies in the virus’s low immunogenicity, which facilitates immune evasion and enables persistent infection^4^. HPV employs multiple mechanisms to induce local immune tolerance in cervical epithelial cells, disrupting immune homeostasis and allowing viral genome integration into host basal epithelial cell nuclei^5^. Subsequent expression of viral oncoproteins, such as E6 and E7, promotes malignant transformation by inactivating tumor suppressor proteins and impairing cell cycle regulation^6^. Despite these insights, the precise molecular and immunological mechanisms underlying immune dysregulation in the cervical tumor microenvironment remain to be elucidated.

In recent years, immunomodulatory therapies have emerged as a promising approach to enhance HPV clearance and facilitate the regression of cervical lesions. Among these, the Nocardia rubra cell wall skeleton (Nr-CWS) has shown remarkable clinical efficacy; nowadays, the Nr-CWS treatment has become well accepted to clear HPV infections for cervical SIL patients. Reported one-year HPV clearance and cervical lesion cure rates reach as high as 88% and 91%, respectively^7^, suggesting its potent role in activating local anti-tumor and anti-viral immunity. However, the mechanisms through which Nr-CWS exerts these effects are still poorly characterized, limiting its optimized application and further development.

To address this knowledge gap, our study employs spatial transcriptomics—a cutting-edge technology that enables high-resolution, spatially resolved gene expression profiling within tissue contexts. We compared cervical tissue samples from patients stratified by treatment response: an effective group exhibiting HPV clearance and lesion resolution, and an ineffective group with persistent HPV infection and lesions. Through differential gene expression and pathway analyses, we aim to delineate the immunomodulatory pathways activated by Nr-CWS, with the goal of providing mechanistic insights that may inform future therapeutic strategies

## RESULTS AND DISCUSSION

### Spatial transcriptomics data reveals distinct microenvironments in Nr-CWS responders and non-responders

To elucidate the mechanisms underlying the differential clinical responses to Nocardia rubra cell wall skeleton (Nr-CWS) therapy, we performed spatial transcriptomic profiling on cervical squamous intraepithelial lesion (SIL) tissues from patients who were later treated with Nr-CWS therapy and exhibited either a favorable (HPV clearance/lesion resolution) or an unfavorable (persistent HPV/lesions) outcome (Figure 1A and Table S1).

**Figure 1.**
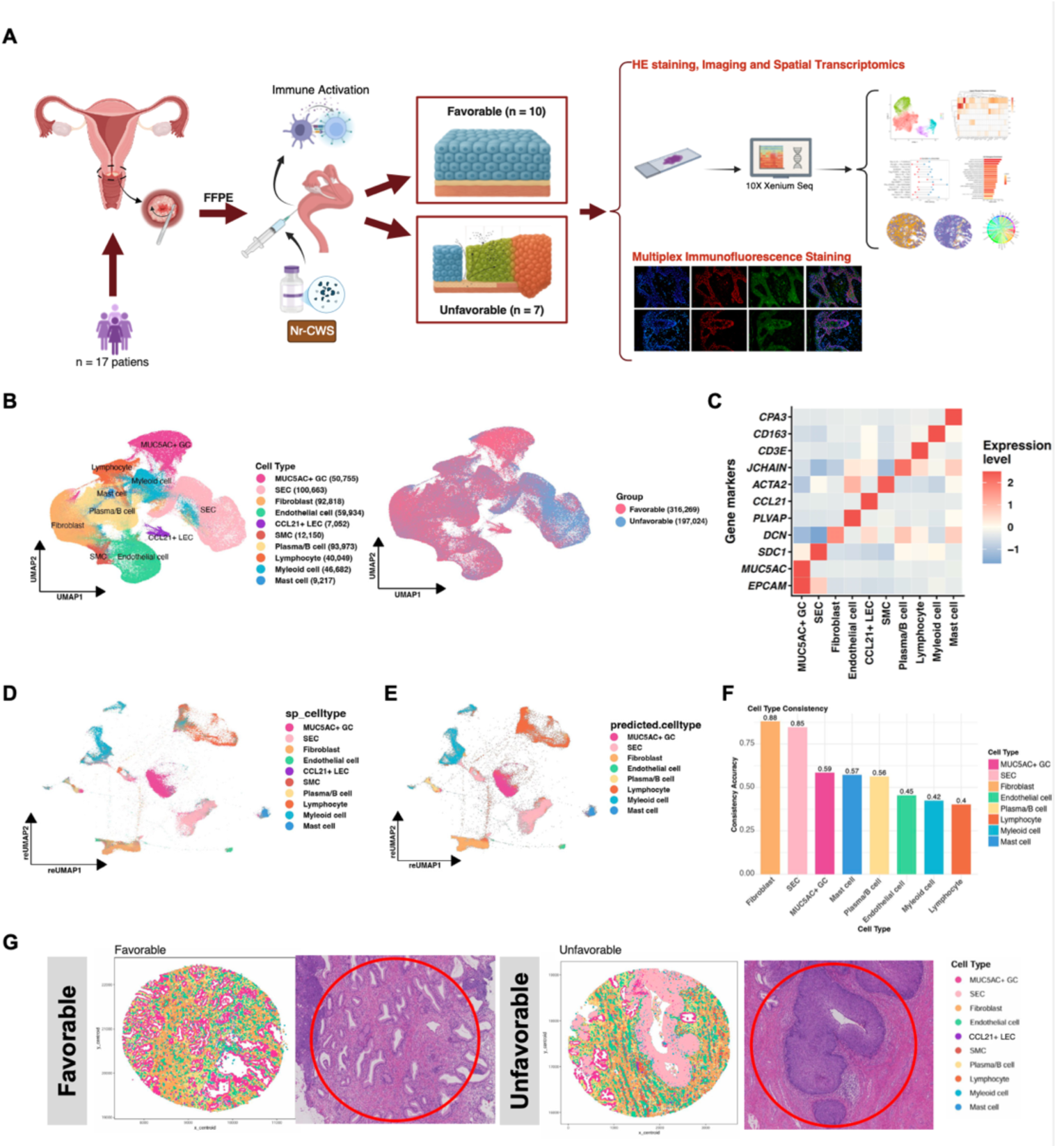
Spatial profiling of different low grade squamous intraepithelial lesion cervical tissues with divergent responses to Nr-CWS. **(A)** Schematic illustration of the experimental workflow. **(B)** Uniform manifold approximation and projection (UMAP) visualizations, showing both the cell type distribution in the dataset and the distribution of cells grouped by clinical status (favorable vs unfavorable). **(C)** Heatmap showing the scaled average expression levels of marker genes across main cell types. **(D)** The UMAP plot indicating the ST data projected into the scRNA-seq UMAP, colors represent the ST major cell type annotations. **(E)** The UMAP plot indicating the ST data projected into the scRNA-seq UMAP, colors represent the predicted major cell type annotations based on scRNA-seq data. **(F)** Bar plot showing the consistency accuracy between original spatial cell type annotations (main_celltype) and scRNA-seq data predicted cell types (predicted.celltype) across shared cell types. **(G)** Two examples showing spatial distributions of major cell types and HE staining images in favorable and unfavorable samples.

We first established a comprehensive cellular atlas of the cervical SIL microenvironment. Uniform Manifold Approximation and Projection (UMAP) analysis revealed distinct clustering of ten major cell types (Figure 1B), which were further annotated based on their canonical marker genes (Figure 1C and Table S2). Using the Xenium spatial transcriptomic (ST) data only, we were able to separate ten major cell types, including MUC5AC-secreting glandular cell (MUC5AC+ GC, with marker genes EPCAM and MUC5AC), squamous epithelial cell (SEC, with marker gene SDC1), fibroblast (with marker gene DCN), endothelial cell (with marker gene PLVAP), CCL21+ lymphatic endothelial cell (CCL21+ LEC, with marker gene CCL21), smooth muscle cell (SMC, with marker gene ACTA2), plasma/B cell (with marker gene JCHAIN), lymphocyte (with marker gene CD3E), myeloid cell (with marker gene CD163) and mast cell (with marker gene CPA3).

To validate the accuracy of our cellular annotations in the SIL tissues with ST data, we integrated our ST data with a single-cell RNA-seq reference^8^. UMAP visualization showed a strong concordance between our original cell type annotations and the cell types predicted by single-cell data mapping (Figures 1D and 1E). This high degree of consistency was quantitatively validated across all shared cell types, confirming the robustness of our initial cellular characterization (Figure 1F). Annotation of CCL21+ LEC was based on the fact that the CCL21 gene was most predominantly expressed in “endothelial cells” in the single-cell RNA-seq reference dataset, and CCL21 expressed in LECs was involved with immune response^9,10^. It’s noteworthy that with the 380 gene panel (Table S3), these CCL21+ LECs were clearly distinguished from other endothelial cells, which wasn’t accomplished by the scRNA-seq data, implying that the Xenium platform has the advantage to identify some cell subtypes using the optimized transcriptome profiles rather than the whole transcriptome profiles.

After the cell type annotation step, we plotted all cells to their original spatial locations, forming the spatial organizations of the cervical tissues for different patients (Figures 1G and S1).

### Distinct cellular compositions and molecular profiles in SIL tissues associated with Nr-CWS treatment response

Initial characterization of the cellular landscape within squamous intraepithelial lesion (SIL) tissues revealed pronounced differences in microenvironmental composition between patients with favorable versus unfavorable responses to Nr-CWS therapy. Comparative analysis of major cell type proportions indicated that the unfavorable response group was characterized by a significant expansion of the SIL squamous epithelial cell (SEC) compartment; in contrast, the favorable response group showed a relative enrichment of reparative fibroblasts and CCL21+ LECs—the latter hypothesized to support robust immune activation and antigen drainage post-treatment (Figure 2A and Table S4).

**Figure 2.**
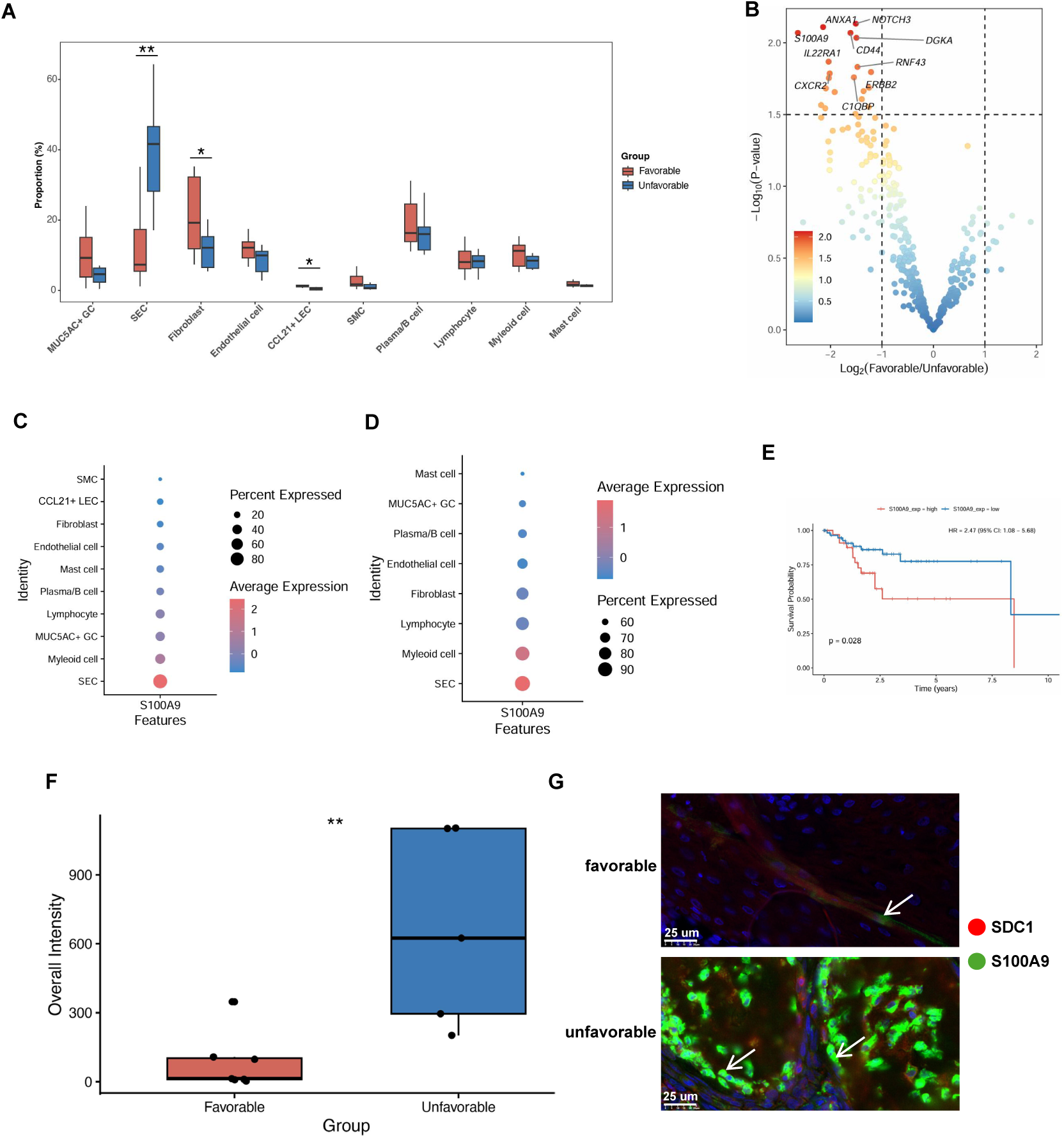
Cellular compositions of the major cell types and bulk gene expression profiles associated with responses to Nr-CWS. **(A)** Boxplot indicating the proportional divergence of major cell types between favorable (red) and unfavorable (blue) clinical groups; statistically significant cell types are highlighted with * (P < 0.05) or ** (P < 0.01) using the student’s t-test. **(B)** Volcano plot showing bulk ST data differential gene expression between Favorable and Unfavorable clinical groups, genes with significant expression changes are shown (p<0.05, |log_2_FC|>1). **(C)** Dot plot showing *S100A9* gene expression across major cell types in spatial transcriptome data (dot color means expression level: blue = low, red = high; dot size indicates percentage of S100A9-positive cells). **(D)** Dot plot showing *S100A9* expression across major cell types in single-cell RNA-seq data (dot color means expression level: blue = low, red = high; dot size indicates percentage of S100A9-positive cells). **(E)** Kaplan-Meier survival curve comparing overall survival between S100A9 high-and low-expression groups in TCGA-CESC squamous cell carcinoma patients (filtered by FIGO stage (IIA–IVB), age ≤70 years, solid tumor tissue, follow-up ≤11 years). **(F)** The bar chart showing S100A9 protein intensity difference between favorable and unfavorable clinical groups; y axis indicates overall intensity normalized by total cell counts in different samples, and statistical result is shown with ** (meaning P < 0.01) using student’s test. **(G)** Two representative examples from favorable and unfavorable groups showing S100A9 immuno-staining results; white arrows highlight coexpression of S100A9 and SDC1.

We next sought to identify overarching transcriptional signatures that distinguish the two clinical groups. Bulk differential gene expression analysis of ST data identified a suite of genes significantly dysregulated in the unfavorable group, offering a high-level perspective on molecular pathways potentially linked to treatment failure. These pathways were notably enriched in inflammatory, cell-cell adhesion and immune evasion-related processes (Figures 2B and S2).

A key finding from this analysis was the consistent and pronounced upregulation of gene S100A9 in the unfavorable group. S100A9 encodes a pro-inflammatory alarmin known to play important roles in chronic inflammation and cancer pathogenesis, often through modulation of myeloid and epithelial cell functions^11–13^. Interrogation of its cellular source within SIL tissues using ST data revealed that S100A9 expression was predominantly localized to squamous epithelial cells, rather than being restricted to classical immune populations (Figure 2C). This unexpected expression pattern was strongly corroborated by an independent single-cell RNA-sequencing dataset derived from two human precancerous lesion samples^8^, which confirmed that SECs serve as the primary expressers of S100A9 in the context of HPV-infected precancerous cervical tissue (Figure 2D). To assess the clinical relevance of this observation, we analyzed data from The Cancer Genome Atlas Cervical Squamous Cell Carcinoma (TCGA-CESC) cohort. Although this cohort represents advanced disease, Kaplan-Meier survival analysis demonstrated that high S100A9 expression was significantly associated with poorer overall survival, supporting its potential role not only in early lesion persistence but also in disease progression (Figure 2E).

Further validation at the protein level via immunohistochemical staining confirmed that S100A9 was substantially more abundant within SECs of the unfavorable group compared to the favorable group (Figures 2F and 2G). This finding reinforces the relevance of epithelial-derived S100A9 in suppressing HPV clearance and potentially facilitating progression toward more severe neoplasia^14^.

In summary, these results define an SIL microenvironment in Nr-CWS non-responders that is typified by an expanded epithelial compartment and a dominant epithelial-inflamed signature centered on S100A9. This signature is not only linked to treatment failure but also portends worse clinical outcomes in cervical squamous lesions.

### Single-cell trajectory analysis reveals dynamic transcriptional changes of cervical precancerous epithelial cells

To investigate the dynamic transcriptional changes underlying cervical precancerous epithelial development and identify key regulators associated with disease progression, we performed pseudotime trajectory analysis on epithelial cells from the single-cell RNA-seq reference. Monocle analysis reconstructed a continuous developmental trajectory, revealing the separation between MUC5AC+ goblet cells (GC) and squamous epithelial cells (SEC) identified in both our ST data and the single-cell RNA-seq data (Figure 3A). This trajectory captured the gradual transition between epithelial lineages, providing a high-resolution view of cellular dynamics during precancerous progression. We then examined the expression pattern of S100A9 along this pseudotemporal continuum, given our previous findings linking epithelial-derived S100A9 to unfavorable Nr-CWS treatment response. Strikingly, S100A9 expression was not uniformly distributed but exhibited much higher expression at the SEC related late pseudotime stages, corresponding to more advanced cancerous states (Figure 3B). This dynamic expression pattern suggest that S100A9 upregulation was coupled with specific developmental transitions during precancerous epithelial maturation, potentially marking a cellular state associated with treatment resistance.

**Figure 3.**
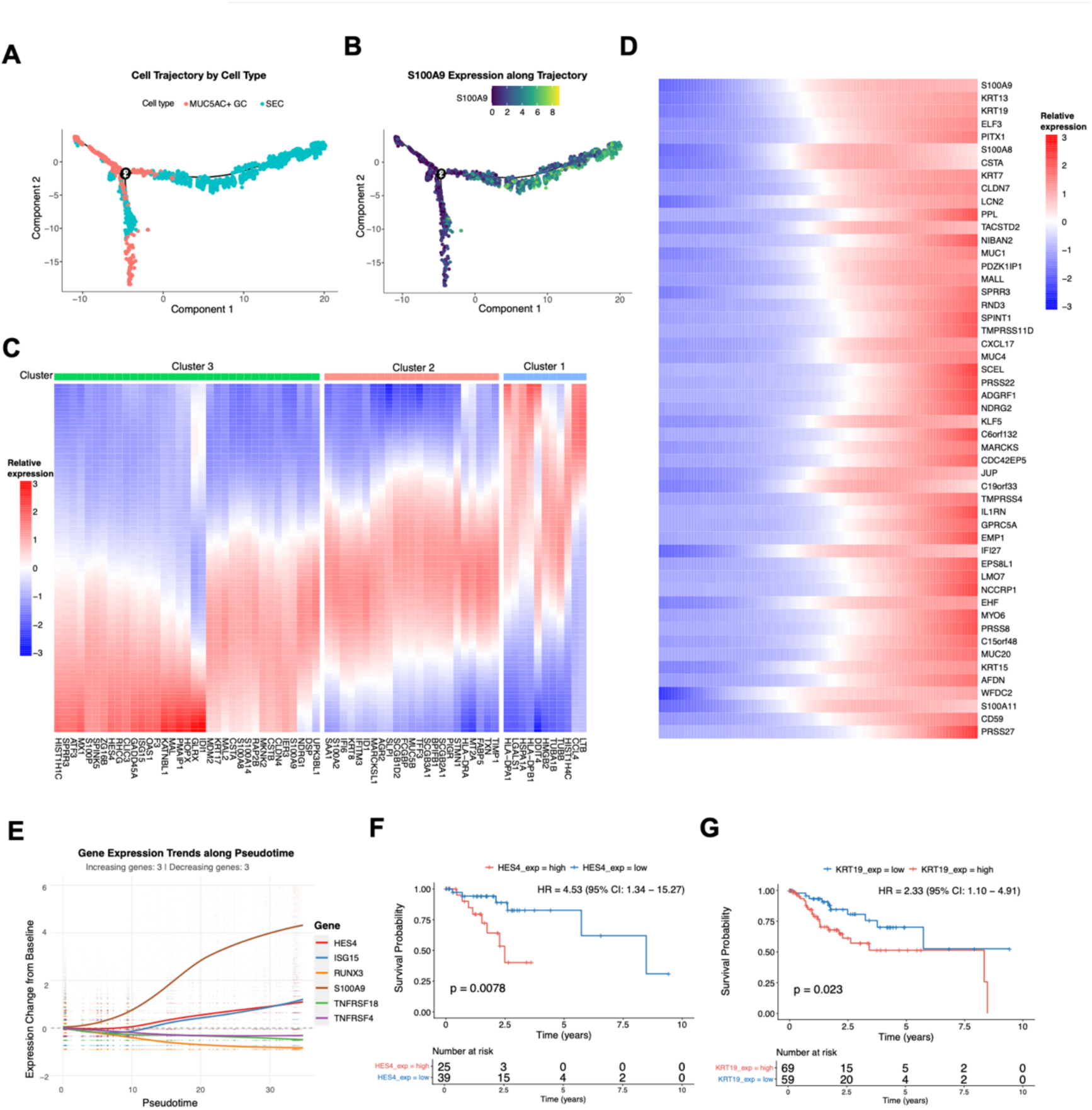
Single-cell trajectory analysis of cervical precancerous epithelial cells. **(A)** Monocle trajectory showing pseudotime trajectory in precancerous cervical epithelial cells and the developmental relationships between MUC5AC+ GC and SEC cells. **(B)** S100A9 gene expression pattern along the pseudotime trajectory in precancerous cervical epithelial cells. Color intensity indicates log-transformed expression level, with dark blue indicating higher expression. **(C)** Heatmap illustrating the dynamic expression patterns of the top 100 pseudotime-associated genes in cervical precancerous epithelial cells. Genes were selected based on the following criteria: q-value < 0.01, expression frequency *≥* 28%, and coefficient of variation (CV) *≥* the 27th percentile. Rows represent individual epithelial cells ordered along the pseudotime trajectory (from early to late), and columns represent genes. Expression levels are depicted using column-scaled values, with red indicating higher expression and blue indicating lower expression. Genes are clustered into three distinct groups (color-coded at the top), each exhibiting unique temporal expression dynamics during epithelial development. **(D)** Heatmap showing the expression dynamics of S100A9 and its top 50 most correlated genes, ranked by Spearman correlation coefficient. Rows represent genes, ordered by the similarity of their expression patterns to that of S100A9; columns represent individual cells, arranged along the pseudotime trajectory from early to late (left to right). Color intensity reflects row-scaled expression levels (Z-score), with red indicating high expression and blue indicating low expression. **(E)** Expression trends of representative genes exhibiting either positive or negative correlation with S100A9. Each colored curve represents the loess-smoothed expression trajectory over pseudotime, accompanied by confidence intervals. Expression levels are shown relative to the baseline (dashed line at y = 0). Genes and corresponding colors are as follows: HES4 (red), ISG15 (blue), TNFRSF18 (green), TNFRSF4 (purple), RUNX3 (orange), JUN (yellow), and S100A9 (brown). (**F, G**) Kaplan-Meier survival analysis of overall survival in TCGA-CESC squamous cell carcinoma patients stratified by HES4 (**F**) and KRT19 (**G**) expression levels. Both analyses were restricted to cases meeting the following criteria: FIGO stage (IA1, IA2, IIA, IIA1, IIA2, IB2, IIIA, IVB), age *≤* 80 years, and survival time *≤* 12 years.

To systematically characterize the transcriptional dynamics underlying epithelial development, we identified the top 100 pseudotime-associated genes based on stringent criteria (q-value < 0.01, expression frequency ≥ 28%, and coefficient of variation ≥ 27th percentile). Unsupervised clustering of these genes revealed three distinct temporal expression patterns, indicating the presence of coordinated transcriptional programs operating at different phases of epithelial carcinogenesis (Figure 3C). Cluster 1 genes such as LTB and HLA-DPA/B1 were highly expressed in early pseudotime and progressively downregulated. Cluster 2 genes showed peak expression at intermediate pseudotime, marking a transitional state characterized by activation of specific signaling pathways. Cluster 3 genes exhibited progressive upregulation toward late pseudotime, including differentiation-associated markers and, notably, S100A9.

Next, we further explored the transcriptional network for S100A9 by identifying the top 50 genes most positively correlated with S100A9 across pseudotime based on Spearman correlation (Figure 3D). Among these genes there were keratin genes such as KRT19, which has been shown to be significantly correlated with HPV-positive oral and oropharyngeal cancers compared to HPV-negative ones^15^; another example was the ETS family transcrioption factor ELF3, which was proven to drive malignant transformation and tumor progression by promoting cell proliferation, survival, and epithelial-mesenchymal interactions^16,17^.

Representative genes exhibiting strong positive or negative correlation with S100A9 were selected for detailed trajectory analysis. Loess-smoothed expression trends confirmed distinct kinetic patterns: HES4 and ISG15 showed positive correlation with S100A9, while RUNX3, TNFRSF18 and TNFRSF4 exhibited negative correlation, with all trajectories displaying well-defined confidence intervals relative to baseline (Figure 3E). The HES4 gene was a member of the Hes family of bHLH transcription factors; it’s overexpression has been observed in several cancer types, and associated with tumor progression, such as in osteosarcoma^18^, colorectal cancer^19^ and hepatocellular carcinoma^20^. Regarding the functions of HES4 in cervical cancer, we performed the Kaplan-Meier survival analysis on TCGA-CESC data to evaluate its clinical relevance. The result demonstrated that high HES4 expression was significantly associated with poor overall survival (Figure 3F). Similarly, we also found high KRT19 expression was correlated with worse patient outcomes (Figure 3G), underscoring the clinical significance of the developmental programs identified in our trajectory analysis.

Together, these findings delineated the dynamic transcriptional landscape of cervical precancerous epithelial cells, identified S100A9-associated gene modules with distinct temporal patterns, and established the prognostic relevance of trajectory-derived genes in cervical cancer.

### Refined sub-cell type analysis uncovers response-associated cellular states

To gain deeper insights into the cellular heterogeneity underlying treatment response, we leveraged higher-resolution Xenium data to subcluster major cell lineages into functionally distinct sub-cell types. UMAP clustering, combined with marker gene heatmaps, successfully identified discrete subpopulations within six major cell classes: squamous epithelial cells (SECs), lymphocytes, MUC5AC+ glandular cells (GCs), myeloid cells, plasma/B cells, and endothelial cells (Figures 4A-F). The remaining four major cell types were not subjected to further subclustering due to either low cell counts or the absence of robust subtype-specific markers in the current gene panel.

**Figure 4.**
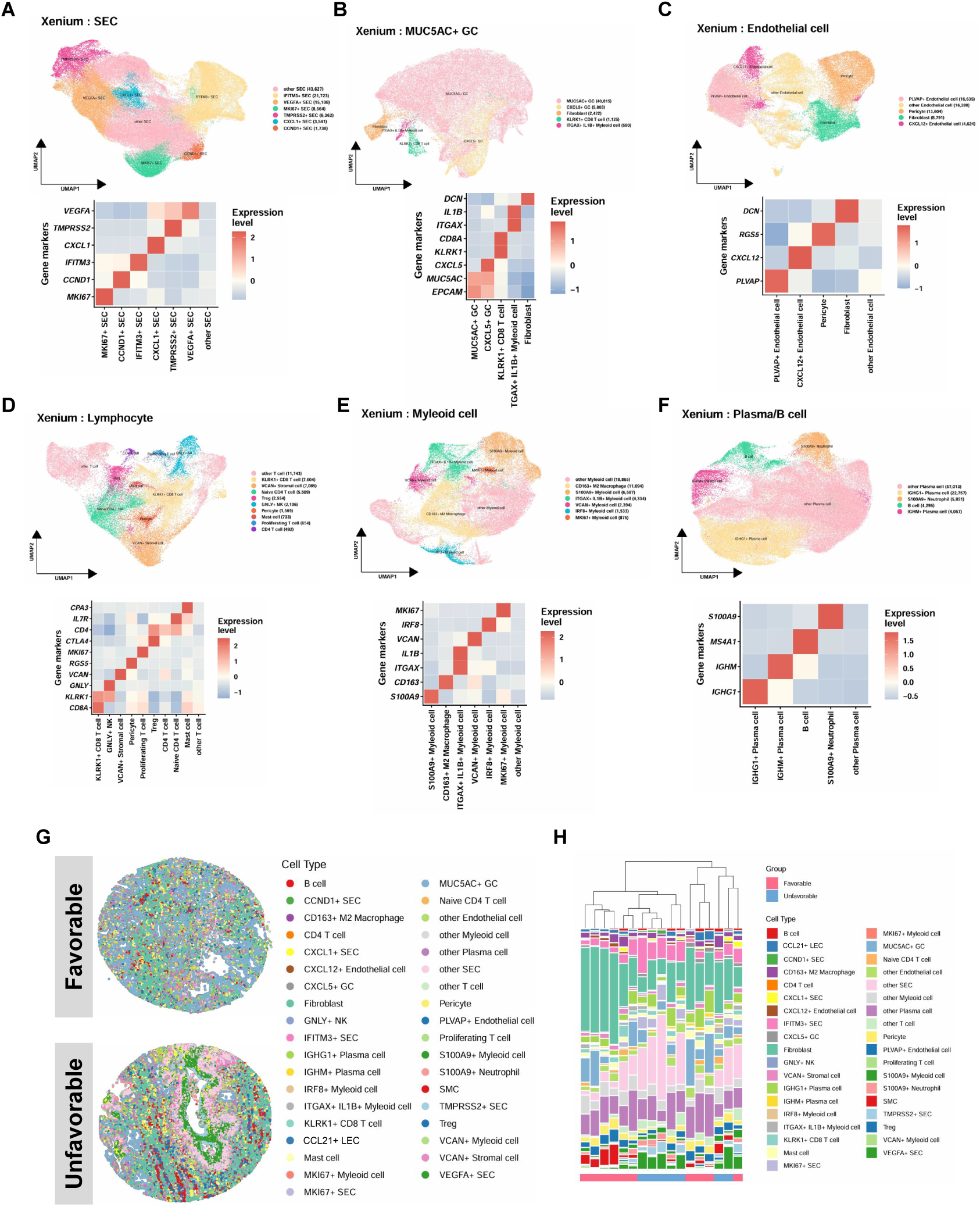
Sub-cell type clustering and annotation of the major cell types based on Xenium Spatial Data. (A-F) UMAP plots (upper) showing sub-cell type clustering for six major cell lineages (SEC, MUC5AC+ GC, Endothelial, Lymphocytes, Myleoid and Plasma/B cell) in Xenium spatial transcriptomic data; the corresponding heatmaps (lower) showing scaled average expression of lineage-specific marker genes across sub-cell types (color intensity indicates expression level: blue = low, cream = mid, red = high). **(G)** Two representative examples for favorable and unfavorable groups showing spatial distribution of sub-cell types. **(H)** Stacked bar chart with hierarchical clustering showing sub-cell type proportions across samples, two colors representing favorable (red) and unfavorable (blue) group samples were shown at the bottom.

We identified six distinct SEC subtypes, each defined by specific functional markers (Figure 4A): IFITM3+ SECs, potentially involved in antiviral defense^21,22^; VEGFA+ SECs, associated with pro-angiogenic functions^23^; MKI67+ SECs, indicative of active proliferation; TMPRSS2+ SECs, with possible roles in cellular entry and signaling^24,25^; CXCL1+ SECs, implicated in neutrophil recruitment and inflammatory responses^26^; CCND1+ SECs, characterized by expression of a key cell cycle regulator previously reported to be upregulated in SIL but downregulated in more advanced lesions^27^. One SEC subset remained unannotated, likely due to the absence of suitable markers in the targeted gene panel.

The MUC5AC+ GC population was further resolved into CXCL5+ GCs, MUC5AC+ GCs, and several non-glandular cell types that were initially co-clustered (Figure 4B). This misclassification may stem from technical limitations such as spillover of MUC5AC signal into adjacent cells during imperfect cell segmentation, or transcriptomic similarities across cell types given the restricted gene panel (380 genes). These non-glandular subtypes included fibroblasts, KLRK1+ CD8 T cells, and ITGAX+ IL1B+ myeloid cells (Figure 4B).

Endothelial cells were subdivided into: PLVAP+ endothelial cells, implicated in vascular permeability and structural integrity^28^; CXCL12+ endothelial cells, which express a pleiotropic chemokine crucial for recruiting CXCR4-expressing immune and progenitor cells^29,30^; other endothelial subtypes, along with pericytes and fibroblasts (Figure 4C).

Lymphocyte subclustering revealed diverse functional states, including naïve CD4 T cells, regulatory T cells (Tregs), conventional CD4 T cells, KLRK1+ CD8 T cells, proliferating CD4/CD8 T cells, GNLY+ NK cells, and additional T cell subsets, as well as minor populations of pericytes, mast cells, and VCAN+ stromal cells (Figure 4D). Myeloid subpopulations were defined by expression of markers such as CD163, S100A9, ITGAX/IL1B, VCAN, IRF8, and MKI67, suggesting diverse roles in inflammation^31,32^, antigen presentation^33^, and immunosuppression (Figure 4E)^31,34^. The plasma/B cell compartment comprised IGHG1+ and IGHM+ plasma cells, alongside a conventional MS4A1+ B cell cluster and a notably mixed subcluster annotated as S100A9+ myeloid cells (Figure 4F).

Spatial visualization and proportion analysis of these sub-cell types revealed profound architectural differences across samples (Figures 4G and S3). However, hierarchical clustering based solely on sub-cell type abundance failed to cleanly separate samples by clinical response (Figure 4H and Table S5). This suggest that cellular composition alone was insufficient to predict treatment outcome, further underscored the potential importance of spatial organization and local cell–cell interactions in mediating Nr-CWS efficacy.

### Cellular subsets and their functional states underlie differential treatment response

Building upon the observed major cellular differences, we next sought to dissect the SIL microenvironment at a higher resolution to identify specific cellular subpopulations and their functional states associated with divergent responses to Nr-CWS therapy.

We first investigated whether proportions of particular cellular subtypes were selectively linked to treatment efficacy. Comparative analysis revealed a starkly divergent cellular architecture between the response groups. The unfavorable response group was characterized by a marked enrichment of two epithelial subpopulations: MKI67+ proliferative SEC and VEGFA+ SEC (Figure 5A). In contrast, the favorable response group showed no significantly enriched subpopulations but exhibited a higher abundance of two major cell types identified earlier: CCL21+ LEC and fibroblast (Figure 2A). These findings suggest that the clinical response to Nr-CWS was underpinned by fundamental shifts in the stromal and epithelial composition of the SIL microenvironment. Spatial mapping further confirmed the precise tissue localization of these cell types and revealed their distinct spatial context, reinforcing the biological relevance of these cellular shifts (Figure S4).

**Figure 5.**
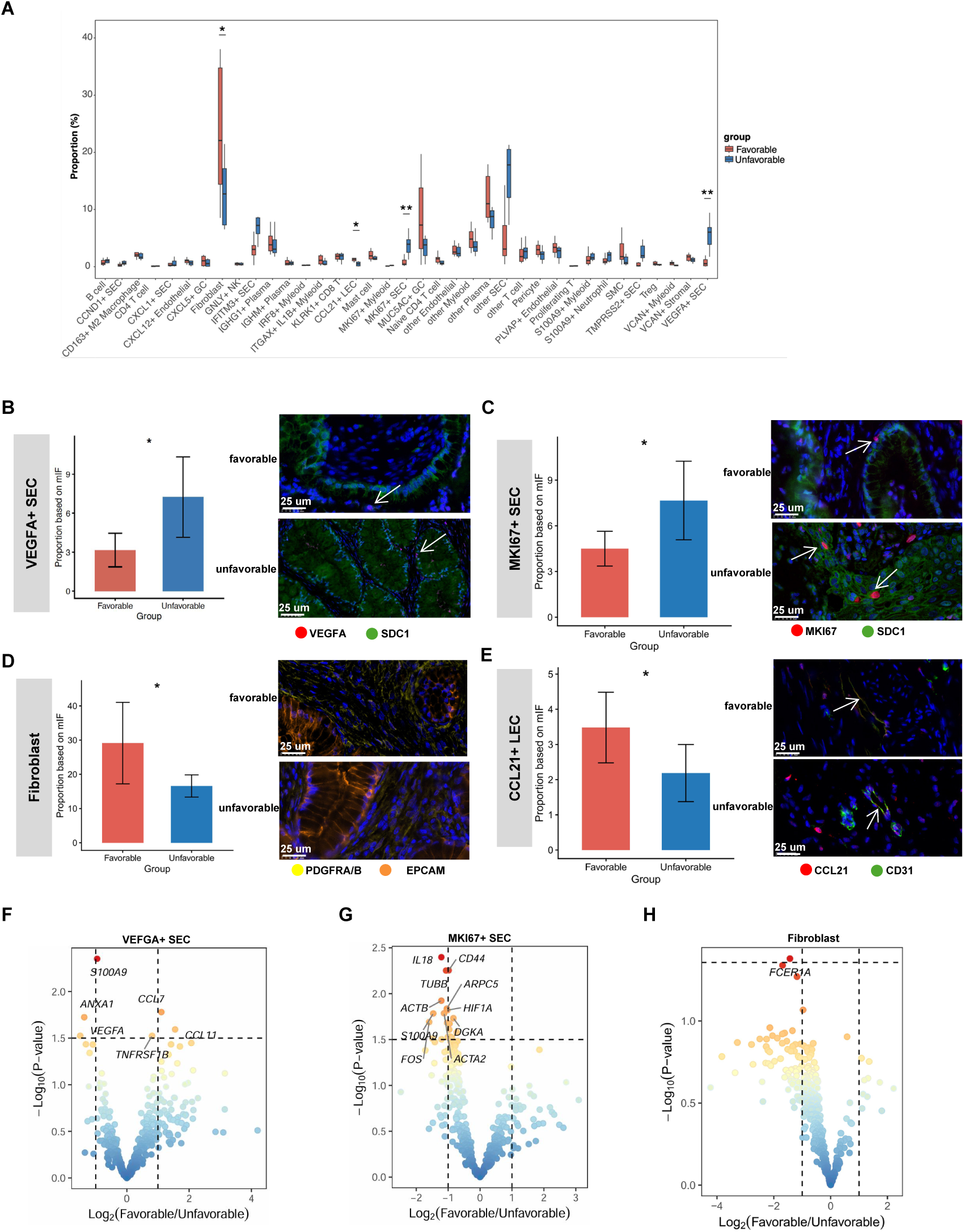
Comparisons of sub-cell type proportions and transcriptional profiles between favorable and unfavorable clinical groups. **(A)** Boxplot indicating the proportions (%) of all sub-cell types as well as four un-subclustered major cell types between favorable (red) and unfavorable (blue) clinical groups; cell types/subtypes with statistically significant difference are highlighted with ** (p<0.01) or * (p<0.05). **(B-E)** The bar chart showing the proportional difference of VEGFA+ SECs (**B**), MKI67+ SECs (**C**), fibroblasts (**D**) and CCL21+ LECs (**E**) between favorable (red) and unfavorable (blue) samples based on the mIF assay; statistical p value is shown using student’s test. Two representative examples from favorable and unfavorable groups are shown beside, white arrows highlight the corresponding cell type/subtype. **(F-H)** The volcano plot showing average gene expression difference of VEGFA+ SECs (**F**), MKI67+ SECs (**G**) and fibroblasts (**H**) between the favorable and unfavorable samples; the x-axis represents the log2 fold change in gene expression between the ’Favorable’ and ’Unfavorable’ groups (Favorable/Unfavorable), while the y-axis corresponds to the -log10-transformed p-value from the Student’s t-test.

The proportional differences of these four critical subtypes were robustly validated at the protein level in an independent cohort of 12 patient samples using multicolor fluorescence staining and spatial mapping. Quantitative analysis confirmed significantly higher densities of VEGFA+ and MKI67+ SECs in unfavorable group samples (Figures 5B and 5C), and conversely, elevated proportions of CCL21+ LECs and fibroblasts in favorable group samples (Figures 5D and 5E).

To elucidate the molecular mechanisms driven by these key subtypes, we performed differential gene expression analysis on VEGFA+ SECs, MKI67+ SECs, and fibroblasts from favorable versus unfavorable groups. For the CCL21+ LEC subset, no significant differentially expressed genes (DEGs) were detected.

In VEGFA+ SECs, we identified a pronounced pro-tumorigenic signature in the unfavorable group, characterized by significant upregulation of S100A9, ANXA1, and VEGFA itself. This coordinated overexpression suggest the establishment of an immunosuppressive, pro-angiogenic, and pro-inflammatory niche^11,35,36^. In contrast, VEGFA+ SECs from the favorable group displayed higher expression of the chemokines CCL7 and CCL11 than the unfavorable group, which were known to promote immune cell recruitment and activation^37,38^ (Figure 5F). This stark transcriptional divergence shed light on the molecular mechanisms of treatment resistance. The MKI67+ SECs from the unfavorable group also exhibited a distinct gene expression profile, with significant upregulation of genes including CD44 and TUBB, which were linked to tumor progression and therapy resistance^39–43^. Interestingly, IL18, a cytokine generally regarded as an immune activator with reported inverse correlations to cervical cancer progression^44^, was also upregulated in the unfavorable context. This suggest that IL18 might contribute to Nr-CWS resistance through non-canonical mechanisms, or its upregulation may be a correlative rather than a causative factor in this specific therapeutic setting (Figure 5G).

Analysis of fibroblasts revealed a significant upregulation of FCER1A in the unfavorable group (Figure 5H). FCER1A encoded the high-affinity IgE receptor (FcεRI), which was implicated in allergic inflammation and fibroblast activation^45^. Its increased expression pointed toward a state of heightened inflammatory activation in fibroblasts associated with poor treatment outcomes.

In summary, our high-resolution analysis reveals that the response to Nr-CWS in SIL was defined not merely by broad immune cell shifts, but by the precise balance of functionally specialized epithelial, endothelial, and stromal subpopulations. The unfavorable outcome was characterized by hyperproliferative and inflammation-driving epithelial subpopulations, whereas the favorable outcome was associated with CCL21 expressing LECs and fibroblasts that may support immune coordination.

### Spatial neighborhood analysis identifies distinct multicellular hubs linked to clinical outcomes

To elucidate how different cell types spatially organize and interact within the cervical precancerous microenvironment, we performed a multi-dimensional analysis of cellular neighborhoods (CNs). This analytical framework enabled us to identify recurrent, spatially coordinated multicellular communities and evaluate their association with Nr-CWS treatment response. The overall distribution of sub-cell types across all identified neighborhoods revealed distinct and reproducible compositional patterns (Figure 6A). For instance, we identified CN-14, a specialized niche predominantly composed of CXCL12+ endothelial cells and pericytes, which may facilitate immune cell recruitment from circulation during HPV infection and potentially promote angiogenesis through stromal signaling^46^. Additionally, we observed several immune-enriched neighborhoods adjacent to CCL21+ LECs, designated as CNs-1, 5, 7, and 11. These LEC-proximal niches could be categorized into two functionally distinct types: one characterized by abundant B cells and plasma cells (CN-1 and CN-5), and another enriched with myeloid and T cell populations (CN-7 and CN-11). The marked heterogeneity in microenvironmental composition surrounding CCL21+ LECs suggests these niches may support different immunological functions, potentially reflecting varied roles in antigen presentation or immune regulation.

**Figure 6.**
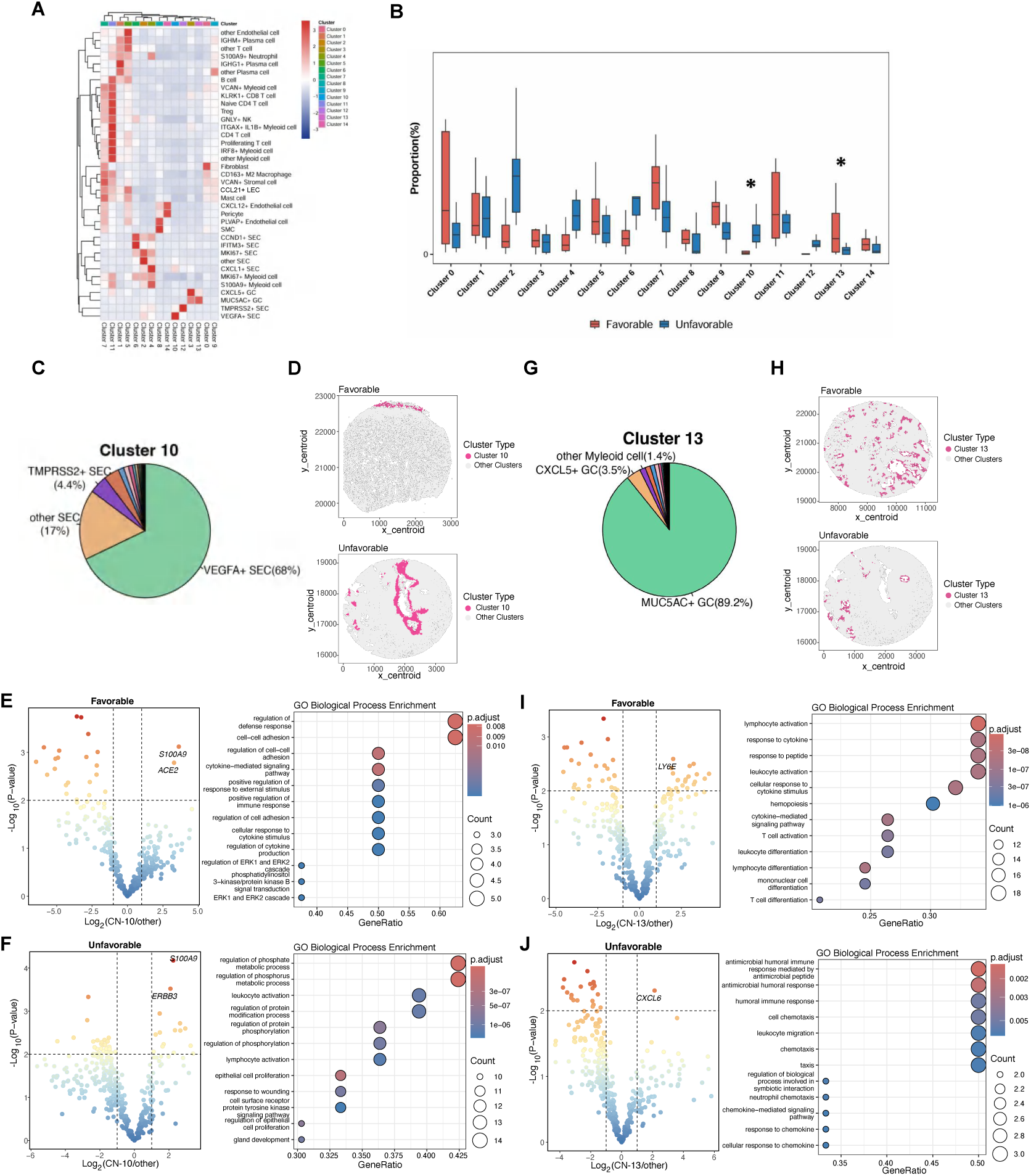
Cellular Neighborhood enrichment analysis in cervical precancerous lesions associated with Nr-CWS responses. **(A)** Heatmap showing the sub-cellular compositions of all 15 cellular neighborhoods in SIL samples; x axis indicates 15 cellular neighborhoods and y axis indicates all cell types/subtypes. Hierarchical clustering was performed on both axis to show similarities. **(B)** Boxplot indicating the proportional difference (normalized by cell numbers) of all CNs between favorable (red) and unfavorable (blue) clinical groups; CNs with statistically significant difference are highlighted with ** (p<0.01) or * (p<0.05). **(C, G)** Pie chart showing the cell type/subtype compositions of CN-10 (**C**) and CN-13 (**G**); different colors indicate different cell types/subtypes, and top three most abundant cell subtypes are highlighted. **(D, H)** Two representative examples from favorable and unfavorable groups showing spatial distributions of CN-10 (**D**) or CN-13 (**H**) in the tissue, highlighted with red colors. **(E, F)** Volcano plots showing bulk differential gene expression analysis result across all cells within CN-10 and other regions in all favorable (**E**, left) and unfavorable (**F**, left) samples separately; the x-axis represents the log2 fold change of average gene expression between the CN-10 and other regions (CN-10/other) and y-axis corresponds to the -log10-transformed p-value from the Student’s t-test. The right panels display the top 12 most significantly enriched Gene Ontology (GO) terms from pathway enrichment analysis of the upregulated genes in CN-10 regions. For the statistical analysis, each sample (rather than each individual CN-10 region) was treated as a replicate to account for inter-sample variability. **(I, J)** Volcano plot showing bulk differential gene expression analysis result across all cells within CN-13 and other regions in all favorable (**I**, left) and unfavorable (**J**, left) samples separately; the x-axis represents the log2 fold change of average gene expression between the CN-13 and other regions (CN-13/other), the y-axis corresponds to the -log10-transformed p-value from the Student’s t-test. The right panels display the top 12 most significantly enriched Gene Ontology (GO) terms from pathway enrichment analysis of the upregulated genes in CN-13 regions. For the statistical analysis, each sample (rather than each individual CN-13 region) was treated as a replicate to account for inter-sample variability.

In addition, comparative analysis of neighborhood prevalence between clinical groups revealed two CNs with particularly strong associations with treatment outcome. Cellular Neighborhood 10 (CN-10) was significantly enriched in the unfavorable response group, whereas CN-13 was predominantly detected in the favorable response group (Figure 6B and Table S6), indicating their potential roles as determinants of therapeutic efficacy.

We further characterized these two critical neighborhoods in detail. CN-10, enriched in unfavorable samples, was primarily composed of VEGFA+ SECs and TMPRSS2+ SECs (Figure 6C), forming dense, localized epithelial hubs as confirmed by spatial visualization (Figures 6D and S5). To decipher the detailed functional states of this neighborhood in response to Nr-CWS treatment, we performed differential gene expression analysis comparing cells within versus outside CN-10. Bulk transcriptomic analysis demonstrated significant upregulation of genes within module CN-10 in samples with favorable outcomes, including S100A9 and ACE2 (Figure 6E); Gene Ontology (GO) enrichment analysis of these upregulated genes revealed activation of defense response and cell-cell adhesion pathways. Conversely, in samples with unfavorable outcomes, the same analysis showed that upregulated genes were primarily enriched for phosphate/phosphorus metabolic processes (Figure 6F); these processes may contribute to Nr-CWS treatment resistance.

In contrast, CN-13 exhibited a markedly distinct cellular composition and spatial architecture. This niche was highly enriched for MUC5AC+ and CXCL5+ glandular cells (GCs) (Figure 6G) and displayed a more dispersed spatial organization (Figures 6H and S6). Transcriptomic profiling of CN-13 regions in favorable samples revealed an immune-activated phenotype characterized by lymphocyte and leukocyte activation, with enrichment of genes such as LY6E (Figure 6I)—a gene known to be involved in modulating viral infection^47^. In unfavorable samples, however, CN-13 niches were predominantly associated with humoral immune response and chemotaxis, marked by enrichment of genes including CXCL6 (Figure 6J). These findings suggest that the functional orientation of the CN-13 niche, rather than its mere presence, may probably influence clinical outcomes after Nr-CWS treatment.

In summary, our analysis demonstrated that the spatial landscape of cervical precancerous lesions was organized into specific multicellular hubs with strong links to therapy response. The unfavorable microenvironment was characterized by CN-10, a pro-proliferative and inflammation-driving epithelial hub, while the favorable microenvironment was defined by CN-13, a mucin and chemokine-producing hub that likely supported mucosal protection and effective local immunity.

### Cell-type-specific spatial microenvironments underlie divergent clinical responses

To dissect the functional impact of specific cellular interactions within the cervical lesion microenvironment, we performed a detailed analysis of the spatial contexts surrounding key “anchor” cell types. This systematic approach revealed that both the cellular composition and molecular programs within a 50 μm radius of these anchor cells were profoundly different between favorable and unfavorable response groups, suggesting that local cellular crosstalk significantly influenced treatment outcomes.

The microenvironment surrounding CCL21+ LECs was overall comparable in the favorable and unfavorable groups. However, a significantly higher proportion of fibroblasts was observed around them in the favorable response group (Figures 7A and 7B). This specific spatial association suggest that interactions between CCL21+ LECs and neighboring fibroblasts may promote lymphatic vessel maturation and function, potentially enhancing antigen drainage and immune cell trafficking following Nr-CWS treatment, possibly through CCL21-CCR7 mediated signaling pathways^48,49^.

**Figure 7.**
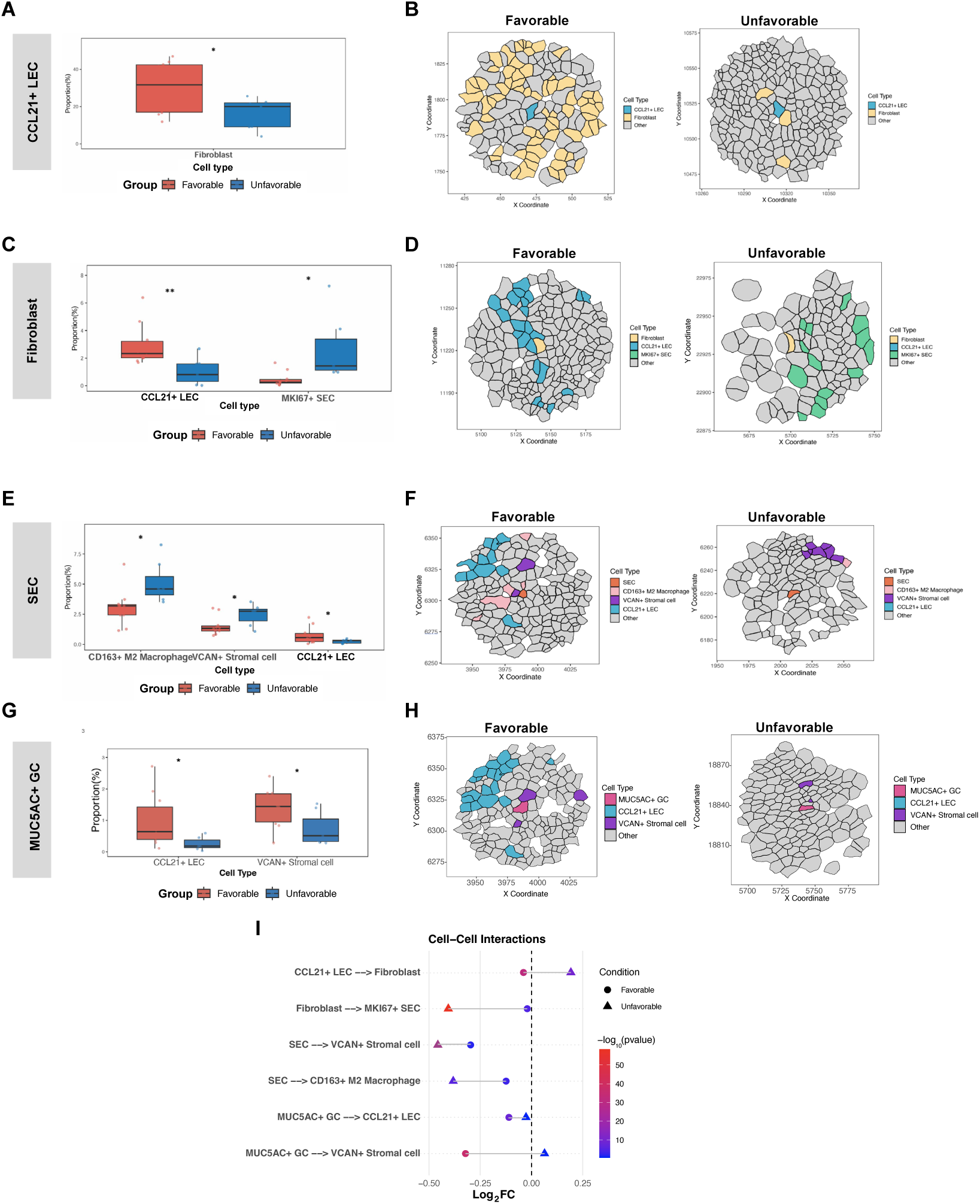
Cell neighborhood frequency analysis in in cervical precancerous lesions associated with Nr-CWS responses. **(A)** Boxplot showing the only one cell type- fibroblast with statistically significant proportional difference between favorable (red) and unfavorable (blue) clinical groups 50 um surrounding the “anchor” CCL21+ LECs; statistically significant cell type-fibroblast is highlighted with * (P < 0.05) using the student’s t-test. **(B)** Two representative examples from favorable (left) and unfavorable (right) groups showing stark proportional difference of fibroblasts 50 um surronding one “anchor” CCL21+ LEC. **(C)** Boxplot showing the cell types/subtypes with statistically significant proportional difference between favorable (red) and unfavorable (blue) clinical groups 50 um surrounding the “anchor” fibroblasts; statistically significant cell types are highlighted with * (P < 0.05) or ** (P < 0.01) using the student’s t-test. **(D)** Two representative examples from favorable (left) and unfavorable (right) groups showing significant proportional difference of CCL21+ LECs and MKI67+ SECs 50 um surrounding one “anchor” fibroblast. **(E)** Boxplot showing the cell types/subtypes with statistically significant proportional difference between favorable (red) and unfavorable (blue) clinical groups 50 um surrounding the “anchor” SECs; statistically significant cell types are highlighted with * (P < 0.05) or ** (P < 0.01) using the student’s t-test. **(F)** Two representative examples from favorable (left) and unfavorable (right) groups showing significant proportional difference of CD163+ M2 macrophages, VCAN+ stromal cells and CCL21+ LECs 50 um surrounding one “anchor” SEC. **(G)** Boxplot showing the cell types/subtypes with statistically significant proportional difference between favorable (red) and unfavorable (blue) clinical groups 50 um surrounding the “anchor” MUC5AC+ GC; statistically significant cell types are highlighted with * (P < 0.05) using the student’s t-test. **(H)** Two representative examples from favorable (left) and unfavorable (right) groups showing significant proportional difference of CCL21+ LECs and VCAN+ stromal cells 50 um surrounding one “anchor” MUC5AC+ GC. **(I)** The dumbbell plots illustrating the interactions between cell pairs as shown in the boxplots above. The X-axis represents the log2-transformed fold change of (actual mean distance/randomized mean distance) for favorable (circle) and unfavorable (triangle) samples. Color intensity indicates the -log10-transformed p-value derived from permutation distance analysis, where red corresponds to lower p-values.

Reciprocal analysis of fibroblast-centered microenvironments further supported this functional relationship. In favorable group samples, fibroblasts were similarly surrounded by a higher proportion of CCL21+ LECs (Figures 7C and 7D), indicating a bidirectional association between these cell types. Conversely, in the unfavorable group, fibroblasts were predominantly encircled by a significantly greater proportion of MKI67+ SECs (Figures 7C and 7D). Given that MKI67+ SECs represented precursors to malignant transformation in squamous intraepithelial lesions, their spatial coupling with fibroblasts suggest the establishment of a pro-tumorigenic niche. This observation aligned well with extensive literature documenting the critical contributions of fibroblast-epithelial interactions in promoting tumor progression through various paracrine signaling mechanisms^50,51^.

Another critical finding emerged from analysis of the SEC microenvironment itself. In the unfavorable group, these epithelial cells existed in niches characterized by a significantly higher density of immunosuppressive cell types, including VCAN+ stromal cells and CD163+ M2 macrophages (Figures 7E and 7F). VCAN (versican) encodes a large extracellular matrix proteoglycan demonstrated to promote tumor progression through multiple mechanisms, including enhanced cancer cell survival, proliferation, migration, and inhibition of immune cell infiltration^32,52,53^. Similarly, CD163+ M2 macrophages have been extensively documented to play pivotal roles in tumor progression by suppressing anti-tumor immunity, promoting angiogenesis, and facilitating tissue remodeling^34,54,55^. On the contrary, a higher proportion of CCL21+ LECs encircling SECs was observed in the favorable group compared to the unfavorable group. This specific spatial configuration was predictive of enhanced immune activation and superior treatment outcomes following Nr-CWS treatment. We next investigated the microenvironment of MUC5AC+ glandular cells (GCs), the defining component of the favorable-enriched neighborhood CN-13. A striking pattern emerged: in responsive samples, these GCs were situated within a niche significantly enriched for CCL21+ LECs and VCAN+ stromal cells (Figures 7G and 7H). The contrasting localization of VCAN+ stromal cells, which either suppress SECs or co-localize with protective GCs, points to a central yet context-dependent role for this stromal population in determining the outcome of Nr-CWS immunotherapy.

To validate these observations, we performed a permutation distance analysis to investigate the spatial interactions of these critical cell pairs. The results confirmed the neighborhood enrichment findings, showing that the observed spatial proximity—such as that of MKI67+ SECs to fibroblasts, VCAN+ stromal cells and CD163+ M2 macrophages to SECs—was statistically significant and not due to random distribution in unfavorable samples (Figure 7I). While in favorable samples, the spatial proximity between CCL21+ LECs and fibroblasts, between MUC5AC+ GCs and CCL21+ LECs/ VCAN+ stromal cells was statistically significant. These findings suggest that the specific interactions between these cell pairs were closely associated with Nr-CWS sensitivity.

Collectively, these cell-type-specific spatial analyses demonstrated that clinical outcome to Nr-CWS therapy was not merely determined by the presence or absence of specific cell types, but was critically influenced by their precise spatial organization and the local molecular dialogues they engaged in. The favorable response microenvironment was characterized by structured fibroblast-endothelial interaction zones that may support immune activation, while the unfavorable response was defined by pro-fibrotic, pro-inflammatory, and immunosuppressive cellular niches that collectively support lesion persistence and progression.

## DISCUSSION

The management of persistent HPV infection and associated cervical lesions remains a significant clinical challenge, with immunomodulatory therapies like Nocardia rubra cell wall skeleton (Nr-CWS) showing promising but variable efficacy. While previous studies have documented the clinical outcomes of Nr-CWS treatment, the underlying cellular and molecular mechanisms determining therapeutic response have remained largely unexplored. Our study leverages cutting-edge spatial transcriptomics to deconstruct the complex microenvironment of squamous intraepithelial lesions (SIL) at unprecedented resolution, revealing a coordinated network of cellular compositions, functional states, and spatial interactions that collectively dictate the fate of Nr-CWS therapy.

Our most fundamental finding challenged the conventional view of epithelial cells as passive targets of HPV infection. Instead, we identified them as active architects of the treatment-resistant microenvironment. This was most strikingly exemplified by the discovery that S100A9, a pro-inflammatory alarmin typically associated with myeloid cells^12,56^, was predominantly expressed by squamous epithelial cells (SECs) in non-responding lesions. The upregulation of S100A9 in SECs created a self-reinforcing loop of chronic inflammation that is fundamentally incompatible with successful immune-mediated clearance of HPV. This epithelial-centric inflammatory signature, characterized by the co-expression of S100A9, ANXA1, and VEGFA in specific SEC subpopulations, defined a novel “immunosuppressive epithelial phenotype” that drove treatment failure. The clinical relevance of this finding was underscored by the significant correlation between high S100A9 expression and poorer overall survival in the TCGA-CESC cohort, positioning S100A9 not merely as a biomarker but as a potential central player in cervical lesion pathogenesis and progression.

The functional diversification of the epithelial compartment further illuminated the mechanisms of treatment resistance. We identified six distinct SEC subtypes, each with specialized roles. The expansion of MKI67+ proliferative SECs and VEGFA+ pro-angiogenic SECs in non-responders created a microenvironment poised for growth and neovascularization. Notably, the transcriptional profile of VEGFA+ SECs in non-responders was not simply pro-angiogenic but profoundly immunosuppressive, highlighting a multifaceted role for this subpopulation in sustaining a treatment-resistant niche. The concurrent upregulation of CD44 and TUBB in MKI67+ SECs further suggest enhanced potential for invasion and therapy resistance, painting a picture of an epithelial compartment that had co-opted multiple pro-tumorigenic pathways to evade immune-mediated destruction.

A pivotal insight from our study was that cellular abundance alone was an insufficient predictor of treatment outcome. While we observed significant differences in the proportions of major cell types and subtypes between responders and non-responders, hierarchical clustering based solely on abundance failed to perfectly segregate the clinical groups. This compelled us to look deeper into the spatial organization of the tissue, where we discovered the true determinants of therapeutic response.

The identification of specific cellular neighborhoods (CNs) provided a spatial framework for understanding Nr-CWS efficacy. CN-10, enriched in non-responders and composed of VEGFA+ and TMPRSS2+ SECs, represented a spatially coordinated “pro-tumorigenic hub.” Within this dense epithelial cluster, cells exhibited a coherent pro-inflammatory and pro-survival gene signature (S100A9, ANXA1, CEACAM1/6), effectively creating isolated islands of resistance within the tissue. The physical concentration of these cells likely facilitated potent paracrine signalling that maintains their malignant phenotype and excluded anti-tumor immune cells.

In stark contrast, CN-13, enriched in responders and dominated by MUC5AC+ and CXCL5+ glandular cells, functioned as a “protective hub.” Its more dispersed spatial distribution and expression of mucosal integrity genes (MUC5AC, TFF3) and chemokines (CCL28) suggested a role in maintaining barrier function and recruiting immune cells. This neighborhood likely represented a microenvironmental niche that was permissive to, and potentially enhanced by, the immune-activating effects of Nr-CWS. The coexistence of these diametrically opposed hubs within the same disease spectrum underscored that the functional output of the SIL microenvironment was not an average of its parts but was driven by the spatial segregation and local dominance of specific cellular communities.

Our analysis of cell-type-specific microenvironments revealed that the functional role of stromal cells was context-dependent, defined by their spatial relationships. In favorable responses, we observed a reciprocal spatial association between CCL21+ LECs and fibroblasts. This specific interaction, occurring within a 50 μm radius, likely facilitated the maturation of functional lymphatic networks, which were critical for antigen presentation and the initiation of adaptive immune responses. This “highway” for immune traffic may be essential for the systemic immune activation required for HPV clearance following Nr-CWS treatment.

Conversely, in unfavorable responses, the stromal function was subverted. Fibroblasts were spatially entangled with MKI67+ proliferative SECs, forming a pro-tumorigenic stromal-epithelial unit. This close association likely activated fibroblasts, driving them towards a cancer-associated fibroblast (CAF) phenotype that supported epithelial proliferation and invasion. Furthermore, the finding that SECs in non-responders were surrounded by a cloak of immunosuppressive cells, including VCAN+ stromal cells and CD163+ M2 macrophages, revealed a sophisticated mechanism of immune evasion. Versican (VCAN) produced by stromal cells can directly inhibit T-cell function, while M2 macrophages were known to suppress cytotoxic responses and promote tissue repair that benefits the lesion. This spatial organization created a physical and chemical barrier that shield the infected and transformed epithelial cells from immune surveillance, effectively negating the immunostimulatory intent of Nr-CWS therapy.

Our findings offer compelling, albeit indirect, insights into the mechanism of action of Nr-CWS. The therapy’s success appear to depend on a pre-existing or inducible microenvironment that was capable of mounting a coordinated immune response. This “permissive” microenvironment was characterized by: 1) a relative lack of the immunosuppressive epithelial phenotype; 2) the presence of protective cellular hubs like CN-13; and 3) structured stromal-immune interactions, particularly between CCL21+ LECs and fibroblasts, that facilitated immune cell trafficking.

In non-responders, Nr-CWS likely failed because it cannot overcome the deeply entrenched immunosuppressive network. The dominant S100A9+ epithelial phenotype, the isolated pro-tumorigenic hubs (CN-10), and the immunosuppressive stromal cloak collectively created a barrier that was insurmountable by the therapy in its current form. This explained why simply activating the immune system was not always sufficient; the local tissue landscape must be capable of receiving and executing the immune-activating signal.

These insights had direct and significant clinical implications. First, the S100A9+ epithelial signature and the spatial metrics (e.g., prevalence of CN-10, fibroblast-SEC proximity) we identified could be developed into powerful predictive biomarkers to stratify patients prior to treatment, sparing non-responders an ineffective therapy and guiding them towards alternative options. Second, our work suggest novel therapeutic combinations. For instance, Nr-CWS could be combined with agents that targeted the S100A9 pathway, disrupt the VCAN-mediated immunosuppression, or reprogramed M2 macrophages to dismantle the resistant niche and sensitized the microenvironment to immunotherapy. The discovery that key resistance mechanisms were operational even at the precancerous (SIL) stage highlight the urgency of understanding and targeting the microenvironment early in the carcinogenic cascade.

Admittedly, this study has several limitations. The 400-gene panel of the Xenium platform, while allowing for high-resolution spatial mapping, necessarily provide a targeted view of the transcriptome. A whole-transcriptome spatial approach in future studies could uncover additional pathways involved in treatment response. Furthermore, the functional inferences regarding cellular crosstalk, while strongly supported by spatial correlation and gene expression data, require validation through in vitro and in vivo experimental models. Techniques such as organoid-immune cell co-cultures derived from patient samples could mechanistically test the roles of specific SEC subtypes and stromal cells in modulating Nr-CWS efficacy. Finally, a longitudinal study tracking microenvironmental changes in the same patients before and after treatment would be invaluable for confirming the causal role of the identified features in therapeutic resistance.

In conclusion, by applying spatial transcriptomics to the context of Nr-CWS treatment for cervical SIL, we have moved beyond a simple catalogue of cellular players to a dynamic model of the disease. We propose that treatment outcome is determined by the balance between two opposing spatial systems: a pro-treatment network featuring immune-permissive glandular hubs and supportive stromal-lymphatic interactions, and a resistance network dominated by immunosuppressive epithelial hubs and pro-tumorigenic stromal-epithelial collaborations. The failure of Nr-CWS in non-responders is not due to a lack of immune activation, but rather the presence of a spatially organized, multi-cellular defence system that actively neutralizes it. This refined understanding of the SIL microenvironment opens new avenues for patient stratification and the development of rational combination therapies to overcome immunotherapy resistance in precancerous lesions.

## Conclusions

Our study deciphered that response to Nr-CWS immunotherapy in cervical precancerous lesions originates from a pre-existing spatial network of epithelial-stromal-immune crosstalk in the lesion microenvironment. These findings create a roadmap for guiding patient stratification and the rational design of combination therapies to overcome resistance.

## ACKNOWLEDGEMENTS

The authors sincerely express gratitude to Dr. Zhiwen Li and Dr. Yunyun Liu for their scientific and technical input. This study was funded by grants from the National Natural Science Foundation of China (NSFC-82301853, to T.F. Long); Sun Yat-sen Memorial Hospital, Sun Yat-sen University, Horizontal Project Fund(7670025006, to T.F. Long).

## AUTHOR CONTRIBUTIONS

T. Long, M. Ding and B. Wei conceived the study and supervised the project. Q. Sun, J. Li and Y. Lan collected the samples and performed most experiments. B. Wei and Y. Mao carried out statistical and bioinformatics analyses. Y. Sun, Y. Lu and X. Li performed the data curation and date preprocessing. J. Ma helped with data curation and pathological analyses. All authors have read and approved the final manuscript.

## STUDY PARTICIPANT DETAILS

### Human

This study was approved by the Ethics Committee of Sun Yat-sen Memorial Hospital of Sun Yat-sen University (Approval No. SYSKY-2025-204-01), and conducted strictly in accordance with the ethical principles outlined in the Declaration of Helsinki in 2024. Prior to participation, all subjects provided written informed consent after being fully informed of the study’s purpose, assured of confidentiality, and advised that participation was voluntary and could be withdrawn at any time without penalty. All patient data were de-identified prior to analysis, as all data were fully anonymized and no individual can be identified. Only the paraffin-embedded samples were used for both spatial transcriptomic assay and the multi-IF assay (Table S1). Relevant clinical information was collected from medical records.

## METHOD DETAILS

### Spatial transcriptomics using Xenium platform

Formalin-fixed paraffin-embedded (FFPE) tissues were sliced into 5 µm thick sections, which were then mounted onto Xenium slides (cat# PN-1000659, 10x Genomics) in strict accordance with the manufacturer’s protocol (CG000578, Revision F, 10x Genomics). Following a 3 h baking step at 42℃, the slides were preserved in 50 ml centrifuge tubes with desiccant added, and stored at room temperature until required for subsequent experimental procedures. Subsequently, tissue sections underwent deparaffinization and de-crosslinking processes as outlined in the “Xenium In Situ for FFPE” guide (CG000580, Revision E, 10x Genomics), followed by further treatment in line with the instructions provided in the “Xenium In Situ Gene Expression” user manual (CG000760, Revision B, 10x Genomics). This entire workflow involved multiple manual experimental steps, specifically Priming Hybridization, RNase Treatment & Polishing, Probe Hybridization, Post Hybridization Wash, Ligation, Amplification, Cell Segmentation Staining, Autofluorescence Quenching, and Nuclei Staining. Once these procedures were finalized, slide processing was carried out referring to the “Xenium Analyzer User Guide” (CG000584, Revision G, 10x Genomics). For the assay, a pre-designed “Human Immuno-Oncology” gene panel (cat# PN-1000654, Table S3) which comprises 380 genes was used.

### Spatial transcriptomic data processing

The raw count matrix generated by the Xenium Onboard Analysis tool (v3.3.0) was preprocessed in R (version 4.4.1) using the Seurat package (version 5.1.0). Only cells with expression of at least 3 genes (min.features ≥ 3) were kept, leading to more than 99% cells kept. Raw count data underwent normalization via the SCTransform function. Following PCA analysis with the RunPCA function, the dataset was further integrated using RPCA to reduce batch effect. Subsequent clustering and UMAP dimensionality reduction were performed to identify major cell types.

For subclustering of each major cell type, the procedure was repeated starting from the SCTransform step. A specific cell subtype was annotated only when its marker gene showed expression at least twice that in other subclusters (Log2FC > 1) and was expressed in over 60% of cells within the cluster; marker genes with too low expression were not taken into consideration since they may represent a small subpopulation in that subcluster. Subclusters that could not be annotated under these criteria were labeled as “other” for subsequent downstream analyses.

### Spatial distribution visualization of cell populations

Spatial mapping based on the centroid coordinates of each cell was performed to visualize the spatial distribution of annotated cell types across different samples. First, a spatial transcriptomic data with annotated cell type information was loaded. For each sample (defined by unique values in “orig.ident”), sample-specific metadata subsets were extracted. Spatial plots were generated using the ggplot2 package in R, where each cell was represented by a point with its centroid coordinates. To preserve spatial proportions, plots were formatted with a fixed coordinate ratio (1:1).

### Sub-cell type abundance-based clustering and visualization

Sub-cell type annotation data were extracted from the spatial transcriptomic data. For each sample, counts of individual sub-cell types were quantified and converted to proportions (normalized by total cell count per sample) to adjust for sample size variations.

Samples were clustered by sub-cell type proportion profiles using Euclidean distance and complete-linkage hierarchical clustering, yielding two distinct sample groups. A stacked bar plot was generated to visualize sub-cell type proportions across samples—each bar representing a sample, with segments denoting sub-cell types.

### Single-cell to spatial mapping validation and consistency analysis

A single-cell expression object with reference cell type annotations and a mapping result object storing spatial prediction outputs were used—including precomputed ref.umap dimensionality reduction coordinates, along with two cell type labels per cell: celltype (reference annotations from single-cell data) and predicted.celltype (spatially mapped predictions). UMAP plots were generated via the Seurat package’s DimPlot function to visualize the distribution of reference and predicted cell types, grouped by celltype (reference) and predicted.celltype (spatial predictions) respectively.

To quantify agreement between the two annotations, shared cell types between celltype and predicted.celltype were first identified. Metadata was filtered to retain only cells with these common types, and factor levels were aligned to prevent comparison errors. Consistency accuracy for each cell type was calculated using the dplyr package for grouped summarization. This combined visual and quantitative analysis validated the reliability of spatial mapping across cell types.

### Comparison of cell type proportions between favorable and unfavorable groups

Cell type proportion comparisons between “Favorable” and “Unfavorable” groups were performed to identify group-specific differences in cellular composition. First, metadata (including sample origin, cell type, and group labels) was split into two group-specific subsets. For each group, cell type counts per sample were quantified, normalized by total cell count per sample, and converted to percentages to account for sample size bias.

Normalized proportion data from both groups were merged into a single dataset via factor conversion for consistent visualization. Unpaired t-tests were conducted for each cell type to compare proportional differences between groups, with p-values categorized by significance levels (*** p < 0.001, ** p < 0.01, * p < 0.05, and ns p ≥ 0.05). Boxplots were generated to visualize normalized cell type proportions across groups.

### Differential gene expression analysis

To identify gene expression differences between “Favorable” and “Unfavorable” groups, RNA expression data from a combined Seurat object was first aggregated by sample origin and group. Aggregated expression values were normalized through dividing each sample-group combination’s expression counts by the total cell count of the corresponding sample, eliminating sample size-driven bias in expression quantification.

Normalized expression data were partitioned into “Favorable” and “Unfavorable” subsets, and group-level mean expression values were calculated for each gene. Log₂ fold change (log₂FC) between groups was computed (with a small pseudo-count of 0.0001 added to avoid division by zero) to quantify expression differences, while unpaired t-tests (with unequal variances) were performed for each gene to assess statistical significance of group differences. P-values were adjusted using the Benjamini-Hochberg (BH) method to control the false discovery rate (FDR), and -log₁₀(adjusted p-value) was calculated for visualization.

A volcano plot was constructed to visualize differential gene expression, with genes categorized as “Significant” if they met the thresholds of adjusted p-value < 0.05 and |log₂FC| > 1. The top 10 most significant genes (by p-value) were labeled to illustrate global gene expression differences between groups.

### TCGA-based survival analysis based on S100A9 expression

Survival analysis was performed using TCGA-CESC (Cervical Squamous Cell Carcinoma and Endocervical Adenocarcinoma) data to investigate the association between S100A9 expression and patient survival outcomes. First, TPM (Transcripts Per Kilobase of exon model per Million mapped reads) expression data were processed: ENSEMBL gene IDs were trimmed, then mapped to gene symbols using the org.Hs.eg.db database. Unmatched genes were filtered out, and expression values for duplicate genes (by symbol) were aggregated by mean. TPM values were converted to log-scale (2^TPM - 1) to normalize expression distribution, and the processed matrix was retained for subsequent analysis.

Clinical metadata from TCGA-CESC was curated to define survival endpoints: survival events (1 = death, 0 = alive) and overall survival (OS) time (converted from days to years, with OS calculated as days to last follow-up for alive patients and days to death for deceased patients).Subgroup analysis was restricted to squamous cell neoplasms (SCC), with samples further filtered to retain only those meeting criteria: FIGO stage (IIA–IVB), age ≤ 70 years, tumor tissue type, and solid tissue specimen type. OS was additionally capped at 11 years to focus on short-to-medium-term survival outcomes.

S100A9 expression levels in the filtered SCC cohort were stratified into “high” (above cohort mean) and “low” (below cohort mean) groups. Survival curves were generated using the Kaplan-Meier method, with statistical significance assessed via log-rank test. A Cox proportional hazards model was constructed to calculate the hazard ratio (HR) and 95% confidence interval (CI) for S100A9 high vs. low expression, with “low” expression set as the reference group to illustrate the association between S100A9 expression and patient survival.

### Neighborhood enrichment analysis with spatial transcriptomic data

The k nearest neighboring cells were selected as the analysis window, within which different cell types and subtypes were quantified to generate feature vectors containing cell count information. Cluster analysis was then performed on these vectors to identify cell neighborhoods with recurrent compositional features, thereby revealing spatial niches enriched with specific cell populations. Through extensive parameter optimization, the optimal number of cell neighborhoods that best reflected the spatial structure and heterogeneity of the tissue was determined. Finally, the most significantly enriched cell type characteristics within each neighborhood were identified. To compare neighborhood proportions between groups, sample-level neighborhood counts were converted to proportions (normalized by total cells per sample). These proportions were stratified by group, and unpaired t-tests were performed for each neighborhood to assess statistical differences between “Favorable” and “Unfavorable” groups. Significance labels were assigned to illustrate group-specific differences in neighborhood composition (*** p < 0.001, ** p < 0.01, * p < 0.05, and ns p ≥ 0.05).

### Composition and spatial visualization of cellular neighborhood clusters

Sample contribution to each cellular neighborhood cluster was quantified with a pie chart. Cell neighborhood data were filtered to include only these cluster cells, with sample-specific counts converted to proportions (relative to total cluster cells) and sorted by descending proportion. Cell type composition of the cluster was visualized in another pie chart, with sub-cell type counts converted to proportions and sorted by proportion.

After that, spatial distribution across samples was displayed via spatial point plots. CN labels were reclassified into “Cluster 10/13” and “Other Clusters”, with sample-specific plots generated using cell centroid coordinates. Cell type-specific spatial distribution within Cluster 10/13 was visualized for a target sample by merging spatial transcriptomic and neighborhood data, using the same point style as sample-level plots.

### Quantification of the cellular microenvironment and spatial mapping

Quantification of the cellular microenvironment was performed to quantify the proportion of various cell types surrounding a target cell type and visualize distributions via spatial mapping. First, spatial metadata was processed. For each sample, the proximity of other cell types to the target was analyzed using a radius-based nearest neighbor approach (50 μm). Surrounding cell counts within this radius were tallied for non-target types, converted to percentages (relative to total neighbors per sample) to account for sample size variation, and compared between groups using unpaired t-tests. Boxplots were generated to visualize group differences, with “Favorable” and “Unfavorable” groups distinguished, outliers excluded and significance labels overlaid for clarity. Only cell types with significant differences, sorted by p-value, were included in the boxplots.

### Permutation distance analysis

The permutation-based distance analysis was performed as described before^57,58^. First, one annotated cell type was designated as the “index cell”. Delaunay triangulation was applied within biologically independent tissue regions to establish nearest-neighbor relationships, and the observed mean triangulation distance between the index cell and every other cell type was calculated. To determine whether these observed distances reflected true biological associations, a permutation test was carried out in which cell type labels were randomly shuffled within each region 100 times, while keeping the spatial coordinates intact. This generated a null distribution of expected distances. The average distance per field in each permutation cycle was then compared to the observed distance using a Mann-Whitney U test. For each cell pair, we calculated the fold enrichment of the observed distance relative to the mean of the permuted distances. The entire procedure was repeated iteratively, with each cell type serving as the index cell, providing a comprehensive view of the spatial architecture.

### Multiplex immunofluorescence staining

Sequential multiplex immunofluorescence was performed on FFPE tissue sections using a validated TSA-based kit (PANOVUE, cat# 10004100100) according to the manufacturer’s instructions. Briefly, after deparaffinization and rehydration, slides underwent iterative cycles of heat-induced epitope retrieval, incubation with a primary antibody, HRP-conjugated secondary antibody, and incubation with a fluorophore-conjugated tyramide (Opal dye), followed by microwave-assisted antibody stripping. The process was repeated for each marker in the panel (listed in Table S7). Nuclei were counterstained with DAPI prior to cover slipping.

### Image acquisition and analysis

Multispectral images were acquired using the KFBIO panoramic digital slide scanner (cat# KF-FL-005). Whole slide scans were performed at 20x magnification. The resulting multispectral images were analyzed using the Halo software (version 4.1). Single-cell segmentation and phenotyping were then performed based on nuclear (DAPI) and membrane/cytoplasmic marker expression.

### Statistical analysis

All the results were shown as the means ± SEM. Student’s t test was used for the analysis of two groups. Other statistical analyses were clearly stated in the manuscript.

## Availability of data and materials

The spatial transcriptomic data have been deposited at GSA-OMIX database with accession number OMIX013062. Details of publicly available software used in the study are given in the “METHOD DETAILS” section. No custom code or mathematical algorithm that is deemed central to the conclusions was used.

