## supplementary figures for "High resolution spatial transcriptomics decodes the microenvironmental determinants of response to Nr-CWS therapy in cervical precancerous lesions"

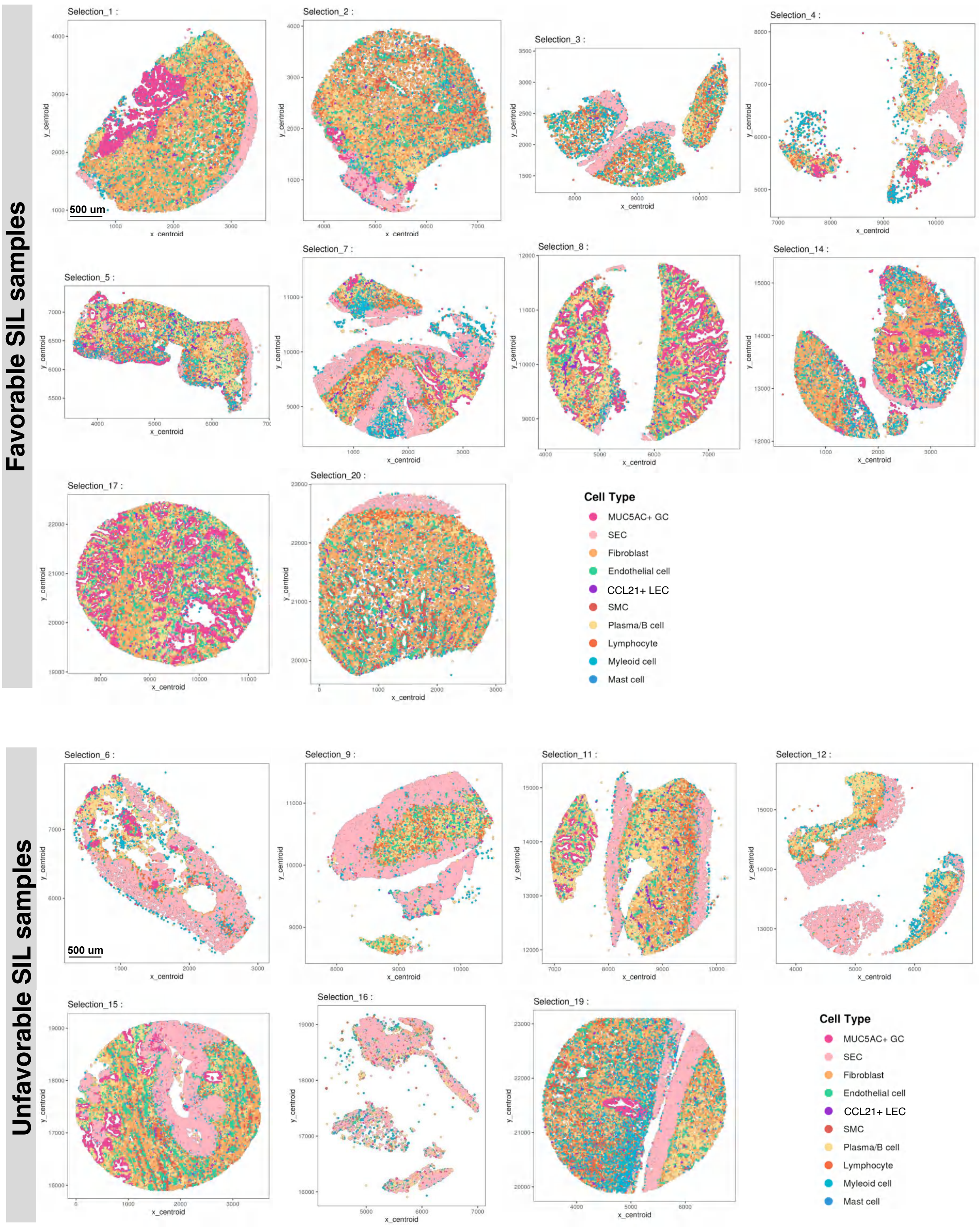


**Figure S1. Spatial visualization of ten major cell types in favorable and unfavorable cervical SIL samples.**

Different colors indicate different major cell types; scale bar is 500 um, which is consistent in all samples, and only shown in the first samples of both groups.

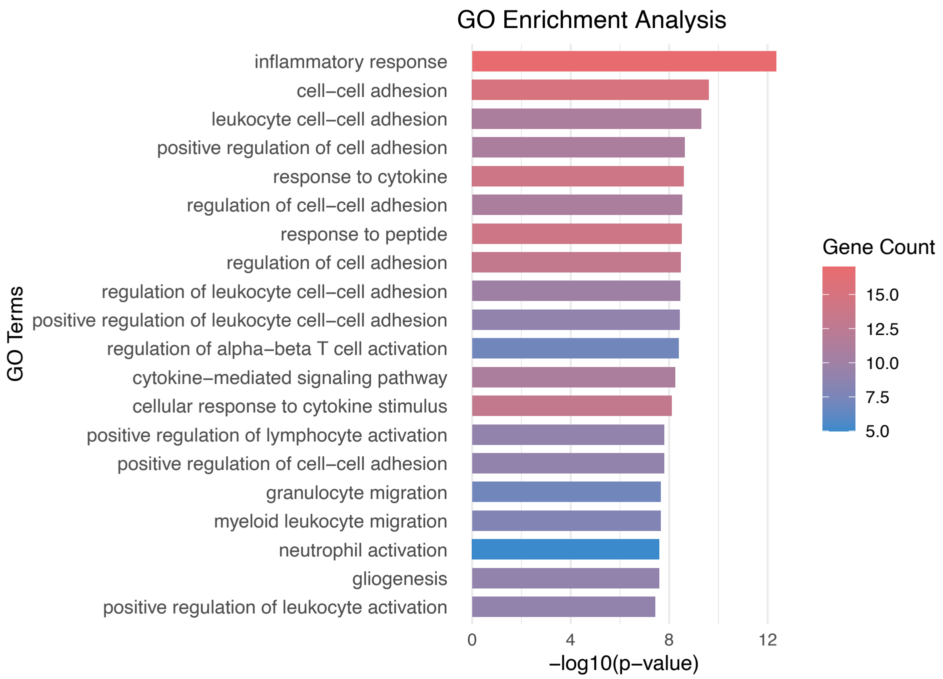


**Figure S2. Gene ontology analysis on the DEGs found in the bulk ST data.**

Different colors indicate different major cell types; scale bar is 500 um, which is consistent in all samples, and only shown in the first samples of both groups.

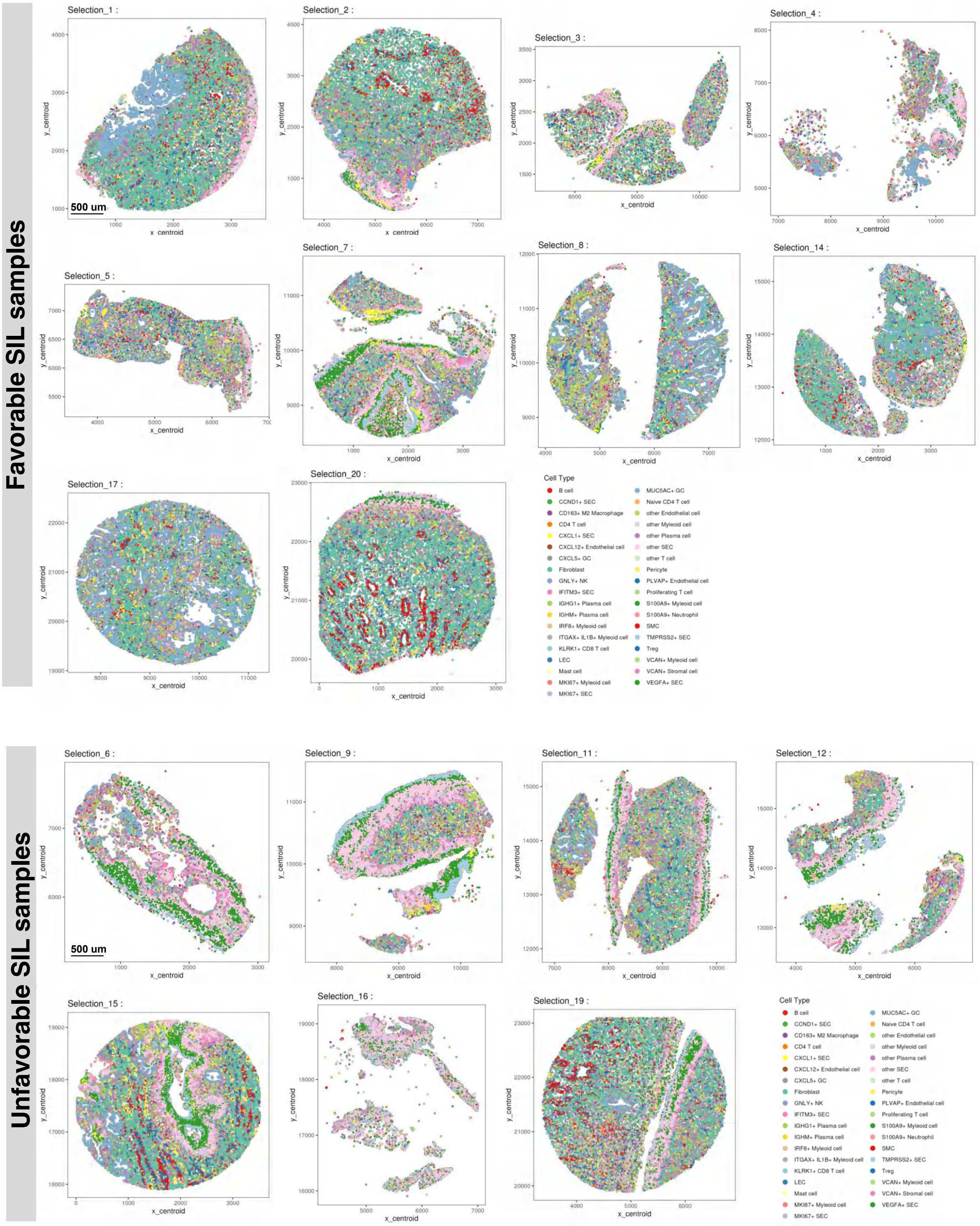


**Figure S3. Spatial visualization of all cell types/subtypes in favorable and unfavorable cervical SIL samples.**

Different colors indicate different major cell subtypes; scale bar is 500 um, which is consistent in all samples, and only shown in the first samples of both groups.

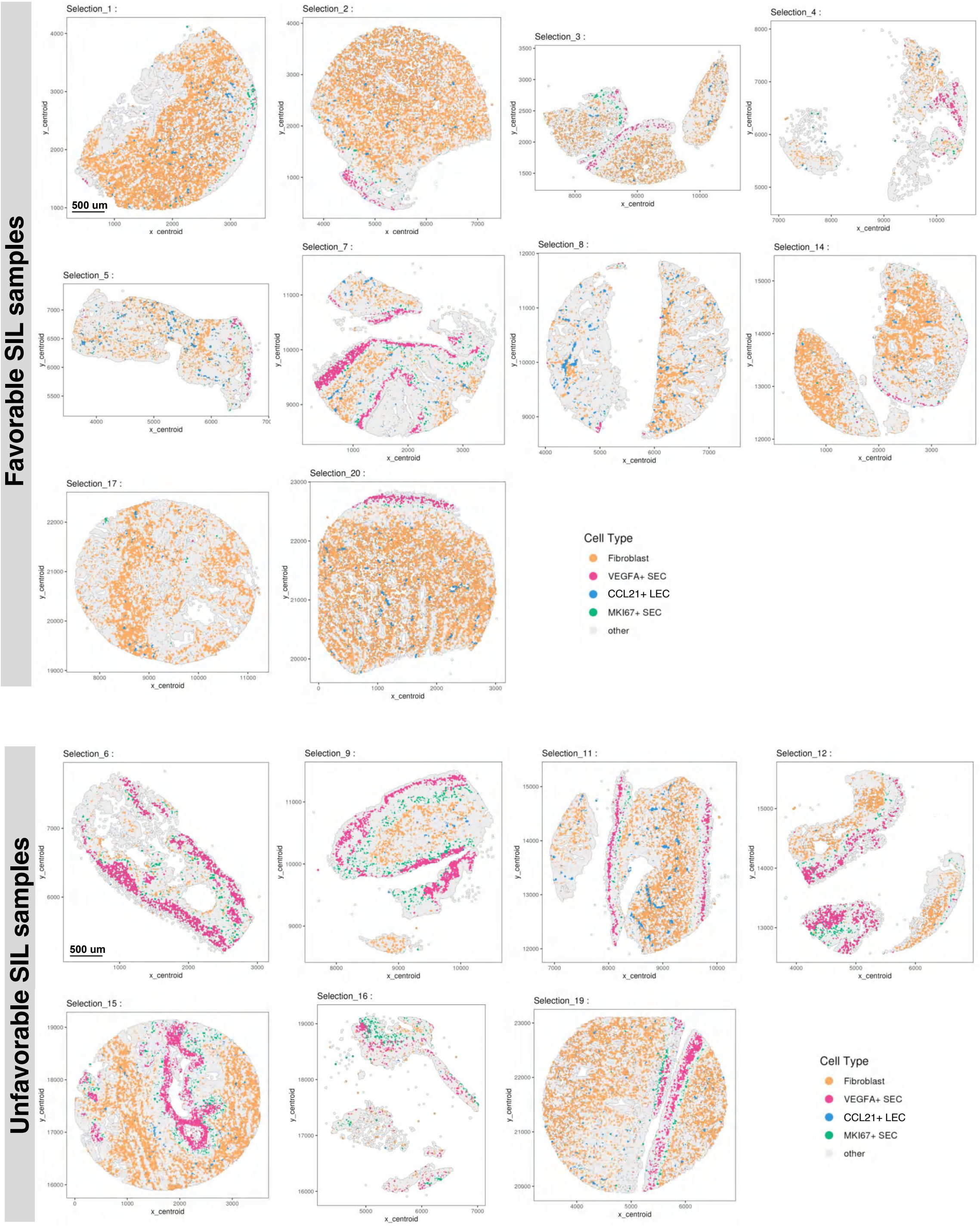


**Figure S4. Spatial distributions of the cell types/subtypes with most significant proportional difference between favorable and unfavorable groups in all cervical SIL samples.**

Different colors indicate different cell subtypes; scale bar is 500 um, which is consistent in all samples, and only shown in the first samples of both groups.

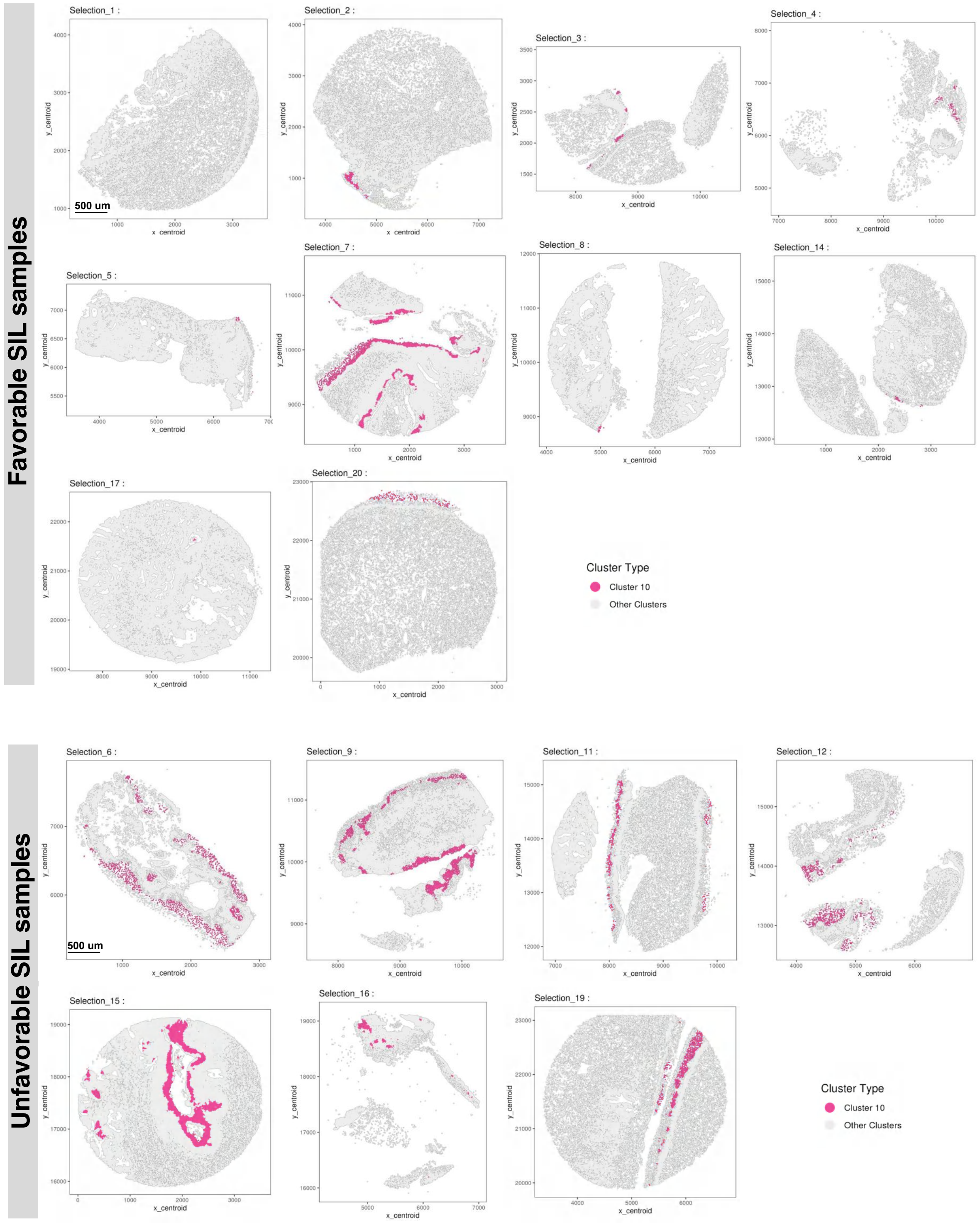


**Figure S5. Spatial distributions of the cellular neighborhood cluster 10 (CN-10) in all cervical SIL samples.**

The red color indicates the CN-10, grey color indicates other clusters; scale bar is 500 um, which is consistent in all samples.

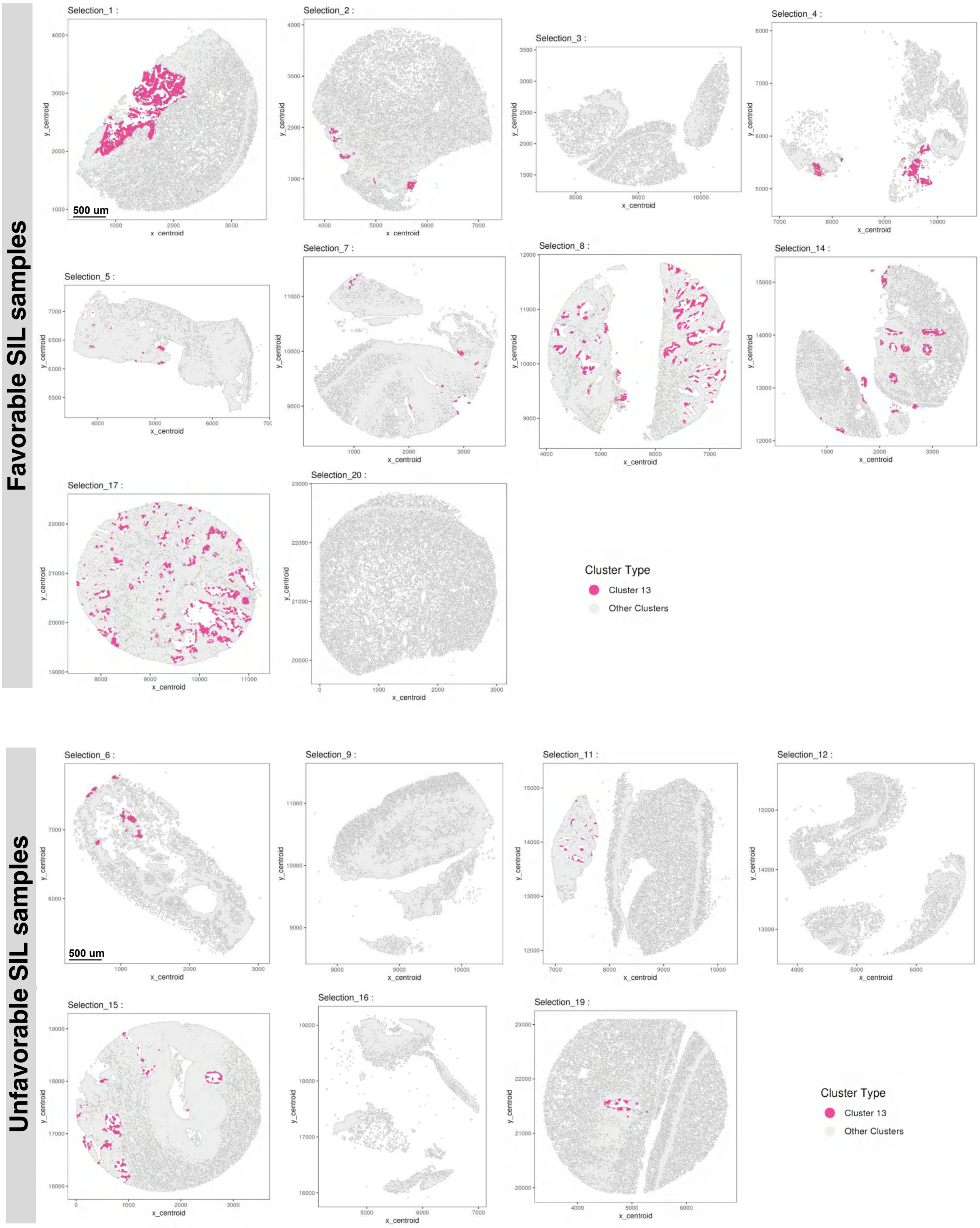


**Figure S6. Spatial distributions of the cellular neighborhood cluster 13 (CN-13) in all cervical SIL samples.**

The red color indicates the CN-13, grey color indicates other clusters; scale bar is 500 um, which is consistent in all samples.
